# The CCL2 Chemokine Promotes Early Seeding of the Latent HIV Reservoir

**DOI:** 10.1101/2021.02.25.432826

**Authors:** Thomas A. Packard, Roland Schwarzer, Eytan Herzig, Deepashri Rao, Xiaoyu Luo, Johanne H. Egedal, Feng Hsiao, Marek Widera, Judd F. Hultquist, Zachary W. Grimmett, Ronald J. Messer, Nevan J. Krogan, Steven G. Deeks, Nadia R. Roan, Ulf Dittmer, Kim J. Hasenkrug, Warner C. Greene

**Affiliations:** J. David Gladstone Institutes, San Francisco, CA 94158 USA; Laboratory of Persistent Viral Diseases, Rocky Mountain Laboratories, National Institute of Allergy and Infectious Diseases, National Institutes of Health, Hamilton, MT 59840 USA; Institute for Translational HIV Research, University Hospital Essen, University of Duisburg-Essen, 45122, Germany; Quantitative Biosciences Institute (QBI) and Department of Cellular and Molecular Pharmacology, University of California San Francisco, San Francisco, CA 94158; Department of Medicine, University of California San Francisco, San Francisco, CA 94110, USA; Department of Urology, University of California San Francisco, San Francisco, CA 94158, USA; Department of Microbiology & Immunology, University of California San Francisco, San Francisco, CA 94158, USA

## Abstract

HIV infects long-lived CD4 memory T cells establishing a latent viral reservoir that necessitates lifelong anti-retroviral therapy (ART). How this reservoir is formed so swiftly remains unknown. We now show the innate inflammatory response to HIV infection results in CCL2 chemokine release, which can drive recruitment of cells expressing the CCR2 receptor including a subset of central memory CD4 T cells. Supporting a role for the CCL2/CCR2 axis in rapid reservoir formation, we find 1) treatment of humanized mice with anti-CCL2 antibodies during HIV infection decreases reservoir seeding and 2) CCR2/5+ cells from the blood of HIV-infected individuals on long term ART contain significantly more provirus than CCR2/5-negative memory or naïve cells. Together, these studies support a model where the host’s innate inflammatory CCL2 response to HIV infection recruits CCR2/5+ central memory CD4 T cells to zones of virus-associated inflammation likely contributing to rapid formation of the latent HIV reservoir.

**GRAPHICAL ABSTRACT:** Why is the latent HIV reservoir established so early following infection? An innate immune response occurs during acute infection that establishes a “zone of inflammation” (step 1). The CCL2 chemokine is produced in part through IFI16 sensing of HIV DNA in abortively infected cells. CCL2 promotes rapid recruitment of CCR2/5+ memory CD4 T cells (step 2). Many of these cells become productively infected (step 3) and a fraction become latently infected (step 4). Thus, HIV hijacks the host inflammatory response to rapidly establish the latent reservoir. In support of this model, we find HIV reservoir reduction in humanized mice treated with anti-CCL2 antibodies during early infection. Further, we find that CCR2/5+ CD4 T cells harbor a substantial fraction of detectable proviruses in the blood of HIV-infected individuals on long-term suppressive ART.

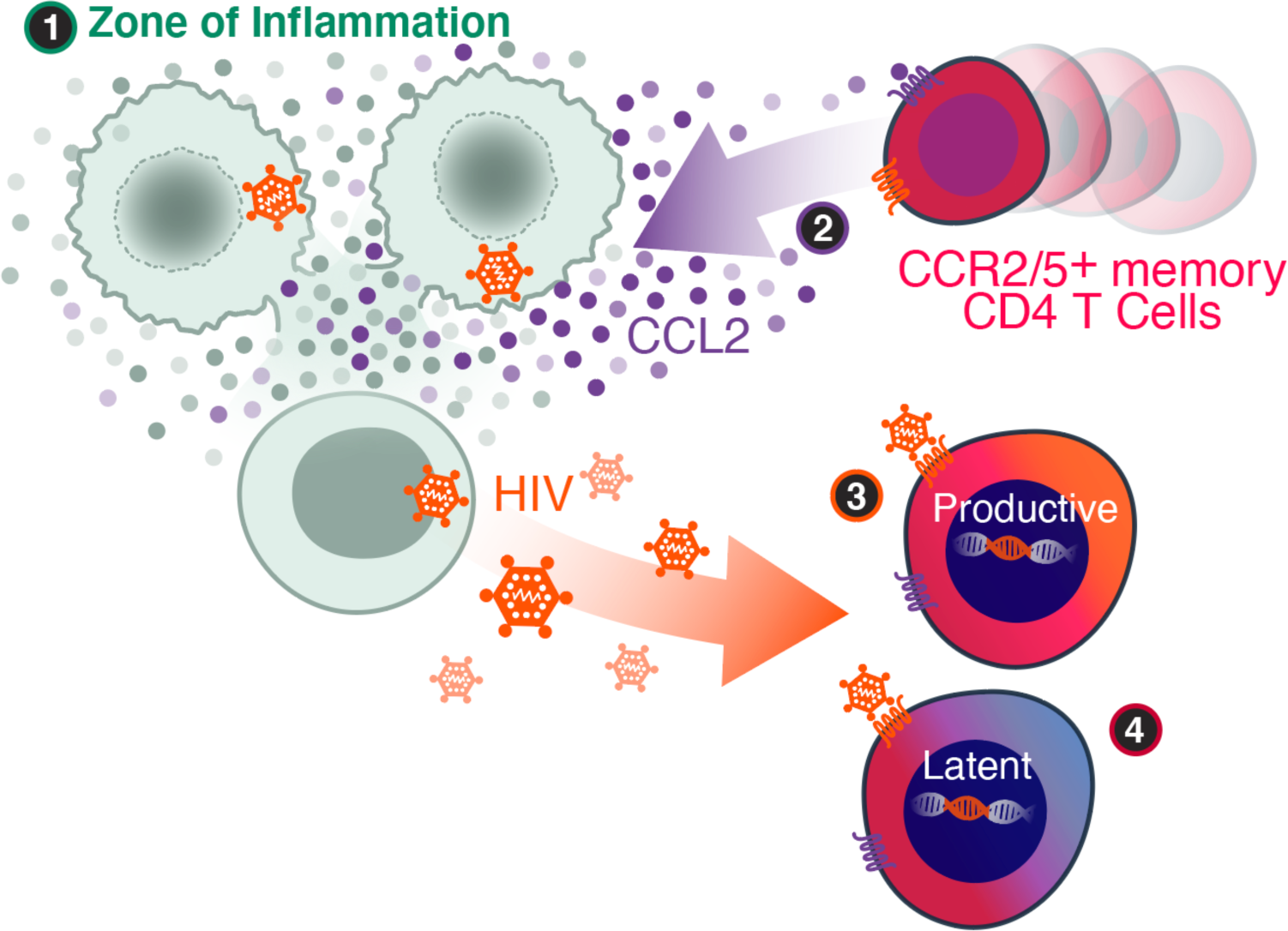

## INTRODUCTION

The latent HIV reservoir is comprised of infectious proviruses predominantly within memory CD4 T cells. These cells are not effectively depleted by antiretroviral therapy (ART), making lifelong treatment a requirement to prevent disease progression. Targeted elimination or control of the latent HIV reservoir is critical for achieving an HIV cure. In rhesus macaques, a stable SIV reservoir is established within 3 days or less following infection ^1^. Likewise, the HIV reservoir is rapidly seeded in humans. Administration of ART immediately after detection of HIV RNA in plasma prior to detection of p24^gag^ or seroconversion (Fiebig stage I) in closely monitored, high risk individuals, fails to prevent viral rebound during a subsequent analytical treatment interruption^2^. Similarly, initiation of ART approximately 10 days after infection (very early Fiebig stage I) does not prevent viral rebound following ART withdrawal ^3^. These findings highlight rapid seeding of the HIV reservoir.

The HIV reservoir is long-lived and self-renewing, as are CD4 central memory T cells. Indeed, many latent proviruses are found within central memory T cells, suggesting they form an important component of the viral reservoir ^4–7^. However, HIV preferentially infects effector CD4 T cells ^8–10^. Effector T cells are terminally differentiated and short-lived. Due to their short lifespan, these cells likely do not contribute to maintenance of the long-lived latent reservoir. How HIV quickly establishes infection in central memory T cells is unknown. HIV may infect central memory precursors, such as stem cell-like memory cells (TSCM) ^11^, although these cells are quite rare. Alternatively, HIV could directly infect central memory T cells.

A potential mechanism for the recruitment of memory cells to the site of infection is via chemoattraction. Acute infection by HIV drives the production of multiple cytokines. Prominent among these is the C-C motif ligand 2 chemokine (CCL2/MCP-1) ^12–14^. CCL2 is one of the first cytokines detected during the acute phase of HIV infection, preceding peak viremia and correlating with viral load ^12, 13, 15^. CCL2 is also involved in the development of the HIV-associated neurocognitive disorder (HAND), attracting cells to cross the blood brain barrier ^16–20^. CCL2’s swift upregulation during acute infection raises the possibility that this chemokine might be involved in the early recruitment of cells involved in latent reservoir formation.

During the course of HIV infection, permissive CD4 T cells can undergo productive infection. Additionally, a large proportion of nonpermissive CD4 T cells in lymphoid tissues are susceptible to abortive infection ^21–23^. These abortively infected CD4 T cells ultimately die by inflammatory pyroptosis ^24, 25^. This response is driven by detection of HIV reverse transcripts by the innate DNA sensor, interferon-inducible protein 16 (IFI16) ^26^. In addition to triggering pyroptosis, IFI16 also activates stimulator of interferon genes (STING). STING then stimulates type 1 interferon (IFN1) via IRF3 phosphorylation ^27^, and CCL2 production via phosphorylation of STAT6 ^28^. We set out to understand the mechanisms that underlie CCL2 production during HIV infection and to characterize the types of CD4 T cells recruited by this chemokine. We describe an intriguing link between innate inflammation, CCL2 production, and the recruitment of CCR2/5+ memory T cells that may facilitate rapid formation of the latent HIV reservoir.

## RESULTS

### HIV triggers IFI16/STING signaling to produce CCL2 in lymphoid CD4 T cells

To investigate whether CCL2 is induced in lymphoid tissue-derived CD4 T cells in response to HIV, we used a previously described *in vitro* cell overlay model ^21^. This method allows for the rapid, synchronous infection of CD4 T cells purified from human tonsils. When measured 18 hours after infection, transcription of CCL2 and IFNB1 were significantly up-regulated while CCL20, an alternative STAT6-regulated chemokine, was not induced (Figure 1A, Figure 1––figure supplement 1). CCL2 protein levels were also increased in the supernatant measured at 24 hours after infection (Figure 1B). In prior studies, blood-derived CD4 T cells exhibited impaired innate immune responses to HIV ^21–23^. Indeed, in contrast to lymphoid tissue CD4 T cells, blood CD4 T cells failed to produce CCL2 (Figure 1––figure supplement 1). Basal levels of CCL2 expression were also increased in uninfected CD4 T cells from lymphoid tissue compared to blood, underscoring the importance of studying tissue-derived T cells in models of HIV infection.

**Figure 1:**
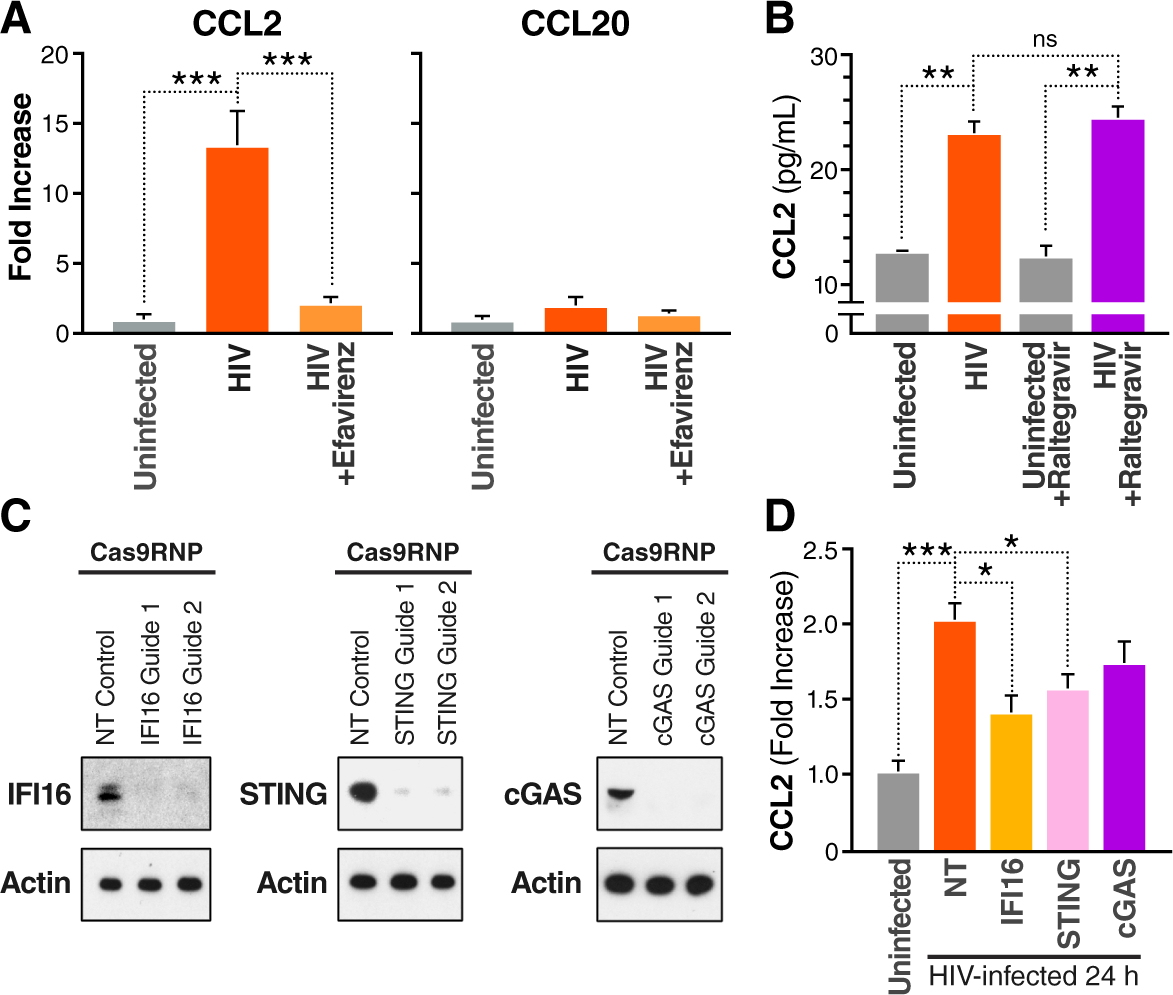
Tonsil CD4 T cells produce CCL2 following HIV infection in part mediated through IFI16 and STING signaling. Tonsil CD4 T cells were purified by negative bead selection and infected with HIV by overlay infection as previously described ^21^. (**A**) Production of CCL2 and CCL20 RNA was measured by qPCR and plotted as fold increase relative to uninfected controls following 18 hours of overlay infection. Treatment with efavirenz (a non-nucleoside reverse transcriptase inhibitor) blocked induction of CCL2. (**B**) CCL2 protein secretion measured by Meso Scale Discovery (MSD) was detected following 24 hours of overlay HIV infection of tonsil CD4 cells. Addition of raltegravir (an HIV integrase inhibitor) did not block production of CCL2. (**C**) Cas9RNPs and two independent guide RNAs (gRNAs) were used to knockout IFI16, STING and cGAS protein expression in primary tonsil CD4 T cells (NT: non-targeting gRNA control). (**D**) CCL2 protein production was significantly diminished (but not eliminated) in HIV-infected cultures when IFI16 or STING were knocked out. Supernatants were harvested 24 hours after infection for MSD analysis of CCL2 production. Fold increase versus uninfected control is shown for each knockout culture. Experiments were performed with 4-6 independent tonsils. Error bars denote SEM; ANOVA with Tukey’s multiple comparisons, * p < 0.05, ** p < 0.01, *** p < 0.001.

To probe which component of the viral lifecycle was required to induce CCL2 production, we employed drugs acting at different steps of HIV replication. CCL2 production was reduced by addition of efavirenz, a non-nucleoside reverse transcriptase inhibitor (Figure 1A), but was unaffected by the integrase inhibitor raltegravir (Figure 1B). Thus, an HIV cDNA intermediate, produced during reverse transcription, appears to trigger rapid CCL2 production by lymphoid CD4 T cells and HIV integration is not required for this response.

This pattern of ART inhibition and lack of response in blood-derived T cells was reminiscent of the pyroptotic cell death response occurring in bystander lymphoid tissue CD4 T cells following abortive HIV infection ^24, 25^. Abortive infection induces pyroptosis through IFI16 sensing of HIV reverse transcripts ^26^. Both IFI16 and the cytosolic DNA sensor cyclic GMP-AMP synthase (cGAS) can activate STING ^27, 29^. In turn, STING can trigger CCL2 production via activation of TBK1 and STAT6 ^28^. To determine which DNA sensors were driving CCL2 production in lymphoid tissue CD4 T cells, CRISPR-Cas9 ribonucleoproteins (Cas9RNPs), were used to knockout the IFI16, STING or cGAS genes (Figure 1C). Knockout of either IFI16 or STING resulted in significantly decreased CCL2 release following HIV infection compared to non-targeted controls (Figure 1D). Knocking out cGAS trended to decrease CCL2 release but did not reach statistical significance. Together, these results indicated that the innate DNA sensing pathway involving IFI16 and STING played an important role in the rapid synthesis and release of CCL2 by lymphoid CD4 T cells in response to HIV infection.

### CCR2 is expressed by a population of CCR5+ central memory cells

A major function of CCL2 production is to recruit cells bearing the CCL2 receptor, CCR2 ^30, 31^. CCR2 is highly expressed on monocytes and is responsible for their recruitment and infiltration within sites of inflammation. Previous studies have shown that CCR2 is also expressed on a unique subset of blood CCR5+ CD4 T cells displaying an effector memory phenotype ^32^.

We set out to examine the phenotype of lymphoid tissue-derived CCR2/5+ cells (Figure 2, Figure 2–––figure supplement 2). The proportion of CCR5+ cells co-expressing CCR2 was substantial: approximately one third of CCR5+ CD4 T cells in the tonsil and two-thirds in the blood (Figure 2D). Using mass cytometry, we compared the expression of 38 markers in blood and lymphoid tissue CD4 T cells (Figure 2––table supplement 1). Using tSNE dimensional reduction visualization, we observed that blood- and lymphoid tissue-derived CD4 T cells sharply differed (Figure 2E). CCR2/5-expressing CD4 T cells between the two compartments also differed, with tonsil-derived cells having elevated expression of CD25 (IL-2Rα), CD7, ICOS, and CCR7 (Figure 2F). CD25 expression is upregulated on activated T cells but constitutively expressed on regulatory T cells. However, the lymphoid CCR2/5+ CD25+ cells did not express FoxP3 so they had an activated rather than regulatory phenotype (Figure 2––figure supplement 3). The high expression of CCR7 in the lymphoid CCR2/5+ cells suggested that these cells were primarily central memory T cells ^33^. In contrast, CD127 and CCR6 were elevated in the blood-derived CCR2/5+ T cells indicating an effector memory phenotype. Together, these findings underscore fundamental phenotypic differences in lymphoid tissue versus blood-derived CCR2/5+ T cells, highlighting the enrichment of central memory in lymphoid tissues, and effector memory in the circulation.

**Figure 2:**
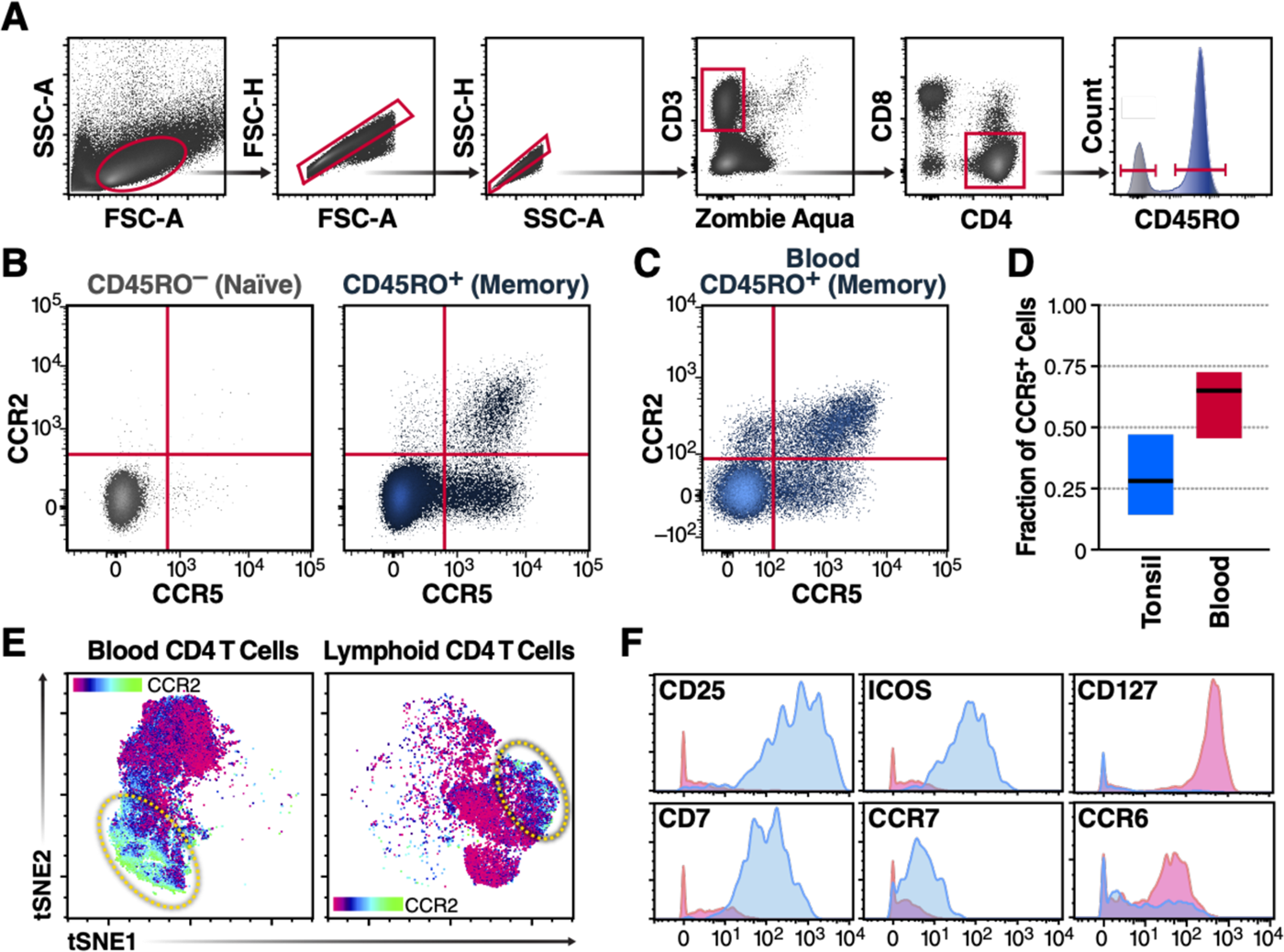
CCR2/5+ CD4 T cells in the lymphoid tissue exhibit a distinct phenotype compared to their cellular counterparts circulating in blood. Freshly isolated mononuclear cells from tonsil and blood were analyzed by multicolor flow cytometry. (**A**) Gating strategy for tonsil memory CD4 T cells is presented. (**B**) CCR2+ expression is primarily limited to CCR5+ memory T cells in tonsil (gated as in panel A). (**C**) Representative chemokine receptor expression of blood-derived memory cells. (**D**) CCR2+ is expressed on a significant fraction of CCR5+ memory CD4 T cells: 28% ± 11% in tonsil (n=18) and 65% ± 10% in blood (n=8). (**E**) Mass cytometry was used to compare blood-derived and lymphoid tissue-derived CD4 cells across 38 different parameters, and tSNE dimensional reduction visualization was performed (n=10 for tonsil and blood). Shown are representative tSNE plots, colored by surface CCR2 expression. (**F**) Selected markers highlight differences between lymphoid tissue (blue) and blood-derived (red) CCR2/5+ cells (See Figure 2––figure supplement 2 for further comparisons).

Comparing CCR2/5+ lymphoid tissue CD4 T cells both to their naïve counterparts and the total memory population shed further light on this unique subset of cells (Figure 3A). The CCR2/5+ cells expressed multiple markers of T cell activation—including surface markers previously shown to be enriched in the latent HIV reservoir. In addition to CD25, these markers included PD1, OX40, and HLA-DR (Figure 3B) ^34–37^. Additionally, these CCR2/5+ cells expressed ICAM (CD54) and LFA-1a (CD11a) which are important in forming the viral synapse ^38^. Finally, they expressed a pattern of surface integrins that reflected potential mucosal origins, including integrins alpha 4 and beta 1, as well as alpha-4-beta-7 ^39, 40^. Taken together, these analyses indicated that lymphoid CCR2/5+ cells were primarily central memory CD4 T cells expressing markers associated with tissue homing, and HIV permissivity—all consistent with a potential role for these cells in establishing the latent HIV reservoir.

**Figure 3:**
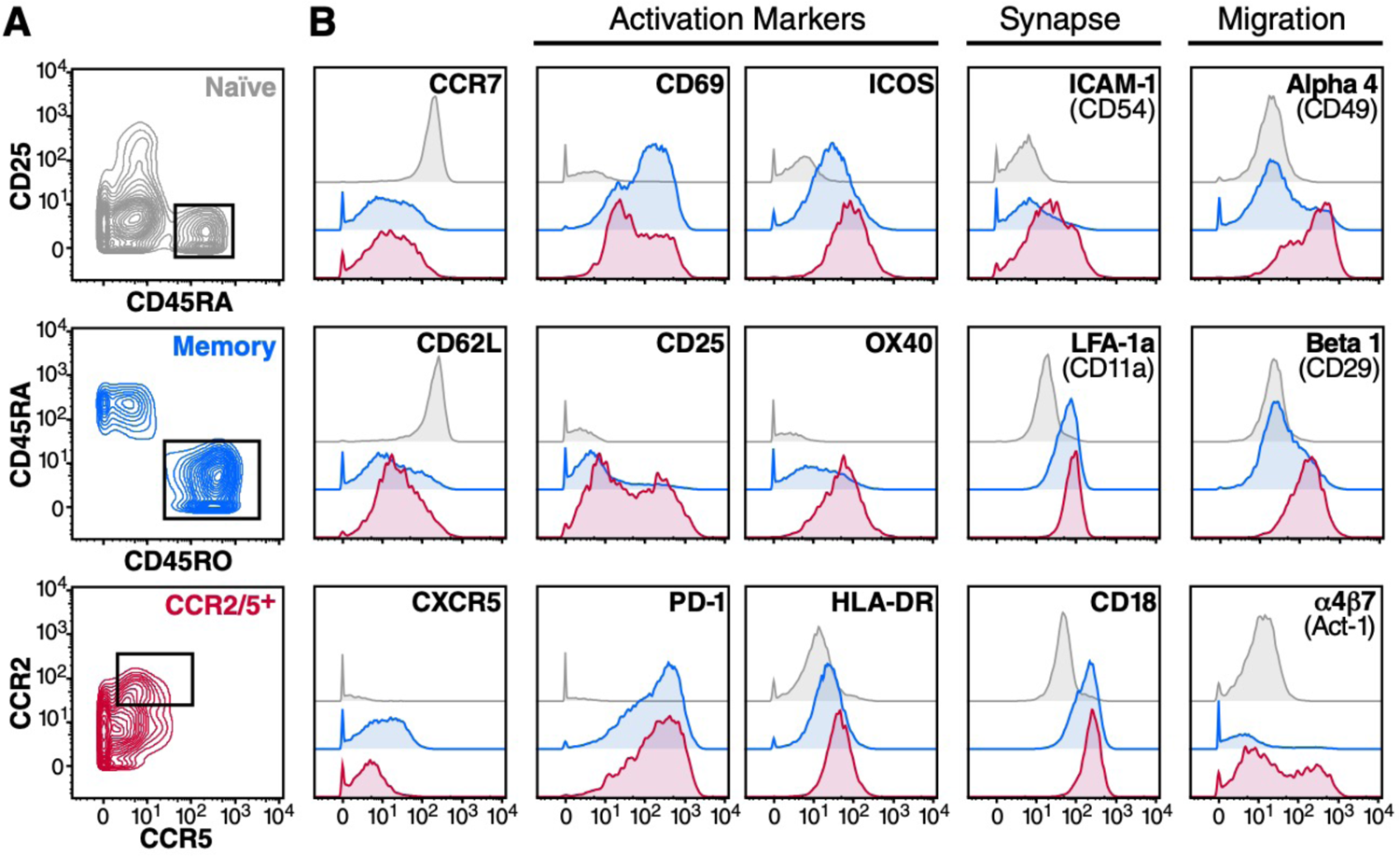
Tonsil CCR2/5+ cells express markers of central memory, activation, viral synapse, and tissue homing. Freshly isolated CD4 T cells from tonsil were analyzed for expression of 38 surface markers by mass cytometry. (**A**) Gating strategy for naïve (pre-gated on CD3+, CD4+, CD8-, CD45RO-, live single cells), memory (pre-gated on CD3+, CD4+, CD8-live single cells), and CCR2/5+ CD4 T cells (pre-gated on memory as above); data shown from representative tonsil (n=10). (**B**) Comparison of marker expression between populations gated as in A (top gray trace, naïve; middle blue trace, memory; bottom pink trace, CCR2/5+).

### CCR2/5+ cells migrate in response to CCL2 and are permissive to HIV infection

It was previously shown that CCR2/5+ CD4 T cells from the blood migrate in response to CCL2 and multiple other chemokines ^32^. Using a transwell migration assay, we found that lymphoid CCR2/5+ cells display a chemotactic response to CCL2 (Figure 4A). CCL2 treatment has been reported to increase the permissivity of blood-derived T cells to HIV infection ^41^. However, we did not observe that exogenous CCL2 affected HIV infection of tonsil-derived CD4 T cells (Figure 4––figure supplement 4).

**Figure 4:**
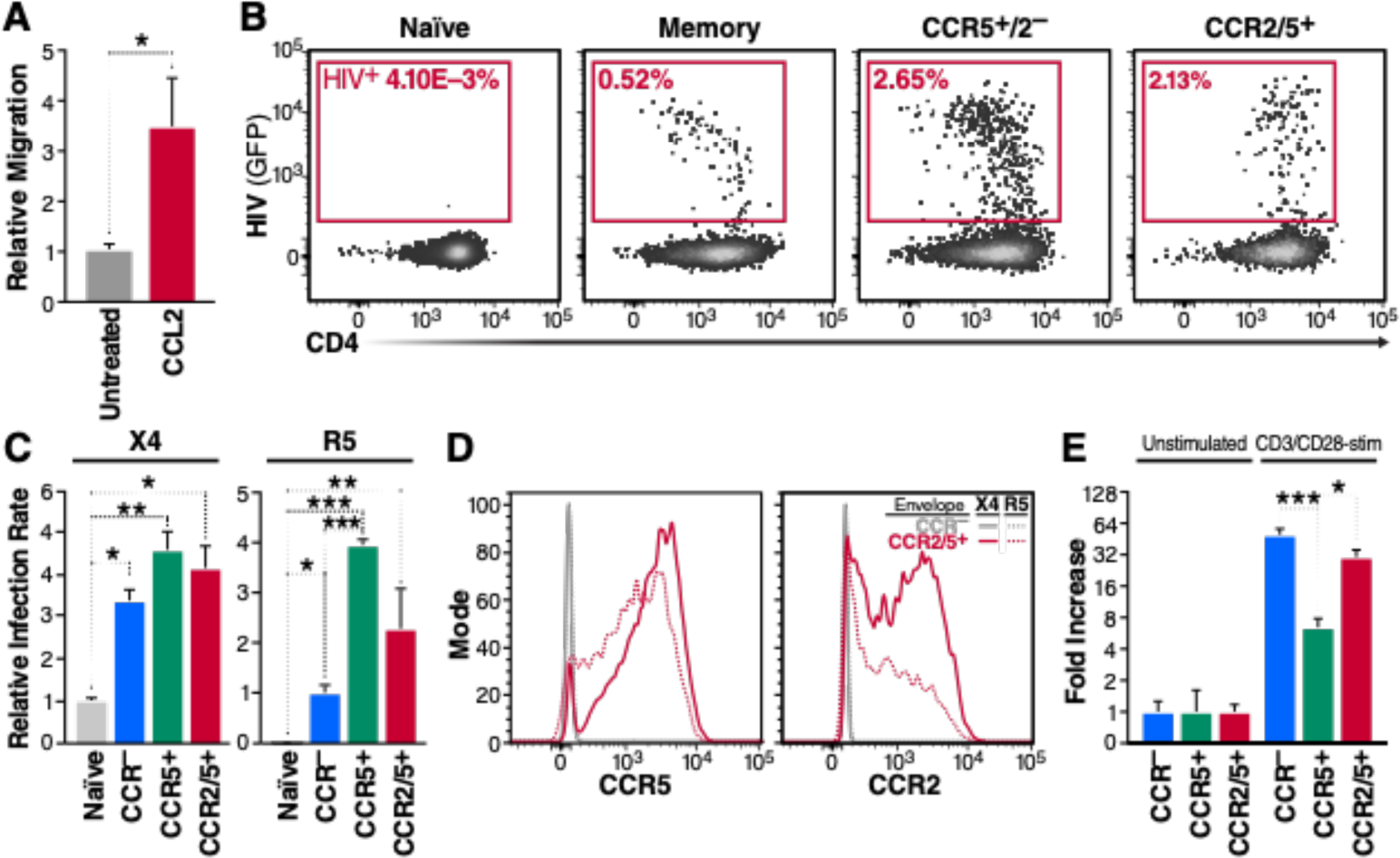
Tonsil CCR2/5+ T cells are permissive to X4- and R5-tropic HIV infection. (**A**) Transwell migration assay of purified tonsil CD4 cells added to the upper chamber of a 2 µm transwell membrane, with 2 ng/mL CCL2 added to the bottom chamber. After 4 hours, the migrating cells (collected from the bottom chamber) were analyzed by flow cytometry and counted. Data are presented as a relative migration (fold increase of chemokine receptor positive over negative cell migration normalized to untreated control). (**B**) Assessment of HIV infection in naïve, memory (chemokine receptor negative), CCR5+, or CCR2/5+ CD4 T cells after sort purification and infection with 100 ng GFP-reporter BaL.NL4-3 HIV-1 (R5-tropic). GFP expression denotes productive infection. (**C**) Relative percentage of GFP-positive cells as gated in panel B, including NL4-3 HIV-1 (X4-tropic) for comparison (n=4). (**D**) R5-tropic envelope causes loss of surface CCR2. CCR5 and CCR2 levels of sorted CCR-negative memory (gray) or CCR2/5+ cells (pink) spinoculated with X4-(solid lines) or R5-tropic (dotted lines) HIV and cultured for 24 hours (**E**) To assess the ability of tonsil CCR2/5+ cells to support latent infection, a previously described primary CD4 T cell latency model was employed ^43^. Cells sorted as in panel B were spinoculated with NL4-3-luciferase reporter virus and cultured in the presence of ART for 5 days. Cells were then reactivated with anti-CD3/28 beads for 24 hours. Virus production after reactivation was measured by quantitating luciferase activity. Data are presented as fold increases in stimulated cells relative to unstimulated cells. Experiments involved the analysis of 4-8 independent tonsil preparations. Error bars denote SEM; ANOVA with Tukey’s multiple comparisons, * p < 0.05, ** p < 0.01, *** p < 0.001.

To assess the intrinsic permissiveness of CCR2/5+ cells to HIV infection, we sorted fresh tonsil T cells and exposed them to GFP reporter viruses expressing either the NL4-3 envelope (X4-tropic) or BaL envelope (R5-tropic). These cells supported productive infection by both viral strains in the absence of *ex vivo* stimulation, (Figure 4B and 4C), demonstrating that in their basal state tonsil CCR2/5+ CD4 T cells were permissive to HIV infection.

Of note, exposure to R5-tropic virus caused rapid loss of surface expression of CCR2 in sorted cells (Figure 4D). This decrease in CCR2 occurred within 24 hours in both infected cells (GFP+) and bystander cells (GFP-) (Figure 4D shows all cells, of which GFP+ are <5%). This effect was not observed following X4-tropic viral infection (Figure 4D, solid lines). A similar downregulation of CCR5 was observed when the R5-tropic envelope was tested. Since this effect was independent of productive infection (observed in GFP-cells), it likely reflects internalization of CCR2 and CCR5 triggered by the binding of the viral envelope. These results highlight the necessity of sorting cells based on their chemokine receptor expression prior to infection, and underscore how exposure to R5-tropic envelope may confound detection and analysis of both CCR5 and CCR2 expression on CD4 T cells.

### CCR2/5+ cells support latent infection by HIV

Only a fraction of tonsillar CD4 T cells (5-10%) undergo productive infection with HIV in the absence of exogenous stimuli. An additional small fraction of cells become latently infected ^42, 43^. To test whether CCR2/5+ cells were able to support latent infection, the cells were sorted as above, then infected with a luciferase reporter virus in the presence of saquinavir (a viral protease inhibitor) to prevent viral spread. The cells were then allowed to rest in the presence of ART to establish latency, as previously described ^43^. After five days in culture, the cells were stimulated for 24 hours with anti-CD3/anti-CD28 antibodies in the presence of the integrase inhibitor, raltegravir, to ensure measurement of post-integration latency. Luciferase expression was measured as a readout of viral reactivation. CCR2/5+ cells underwent similar HIV reactivation after stimulation as CCR2/5-negative memory cells, and significantly greater reactivation than CCR2-/5+ cells (Figure 4E). These findings showed that HIV could establish latent infection in tonsil CCR2/5+ cells *ex vivo*.

We also measured the survival of CCR2/5+ cells following infection with both X4 and R5-tropic virus. We found that the fraction that died following infection was quite similar to that of total tonsil CD4 cells, with approximately two-thirds of cells surviving 48 hours after infection (Figure 4–– figure supplement 5). Together, these results show that HIV infection of CCR2/5+ cells can result in multiple outcomes: productive infection, latent infection, or death.

### CCL2 blockade reduces seeding of the latent HIV reservoir in humanized mice

To understand whether CCL2 played an active role in establishment of the HIV reservoir during acute infection, we used a bone marrow, liver, thymus (BLT)-humanized mouse model of HIV. This model utilized R5-tropic virus allowing the establishment of a latent HIV reservoir in human lymphoid tissue that persists despite ART ^44^. Mice were treated with a blocking anti-CCL2 antibody (n=35) or an isotype control antibody (n=34) 24 hours prior to and during repeated intrarectal HIV inoculations that occurred daily for 5 days (see Figure 5A). The intrarectal infection protocol employed was previously shown to produce ∼50% infection rates ^45^ and was selected to recapitulate mucosal transmission. At day 7 post-infection, mice were switched to ART-infused chow containing tenofovir disproxil-fumerate (Viread), emtricitabine (Emtriva), and raltegravir (Isentress). At day 11 post-infection, anti-CCL2 or control antibody treatment was discontinued. Mice were maintained on ART chow until week 12, and then switched to normal chow. Following treatment interruption, blood was drawn weekly to assess viral rebound. Five animals (14.7%) in the control antibody group exhibited detectable plasma viremia while no animals in the CCL2 blockade group developed detectable viremia (Fisher’s exact p= 0.0248) (Figure 5B). At week 16, the animals were euthanized, and splenic tissue was harvested to measure HIV proviral DNA. In total, 16 animals (47%) from the isotype control arm had detectable HIV, compared to 5 (14%) from the CCL2-antibody treated group (Fisher’s exact p= 0.0041). Together, these findings suggested that treatment with the anti-CCL2-blocking antibody, but not an isotype-control antibody, significantly reduced seeding of the latent HIV reservoir *in vivo*.

**Figure 5:**
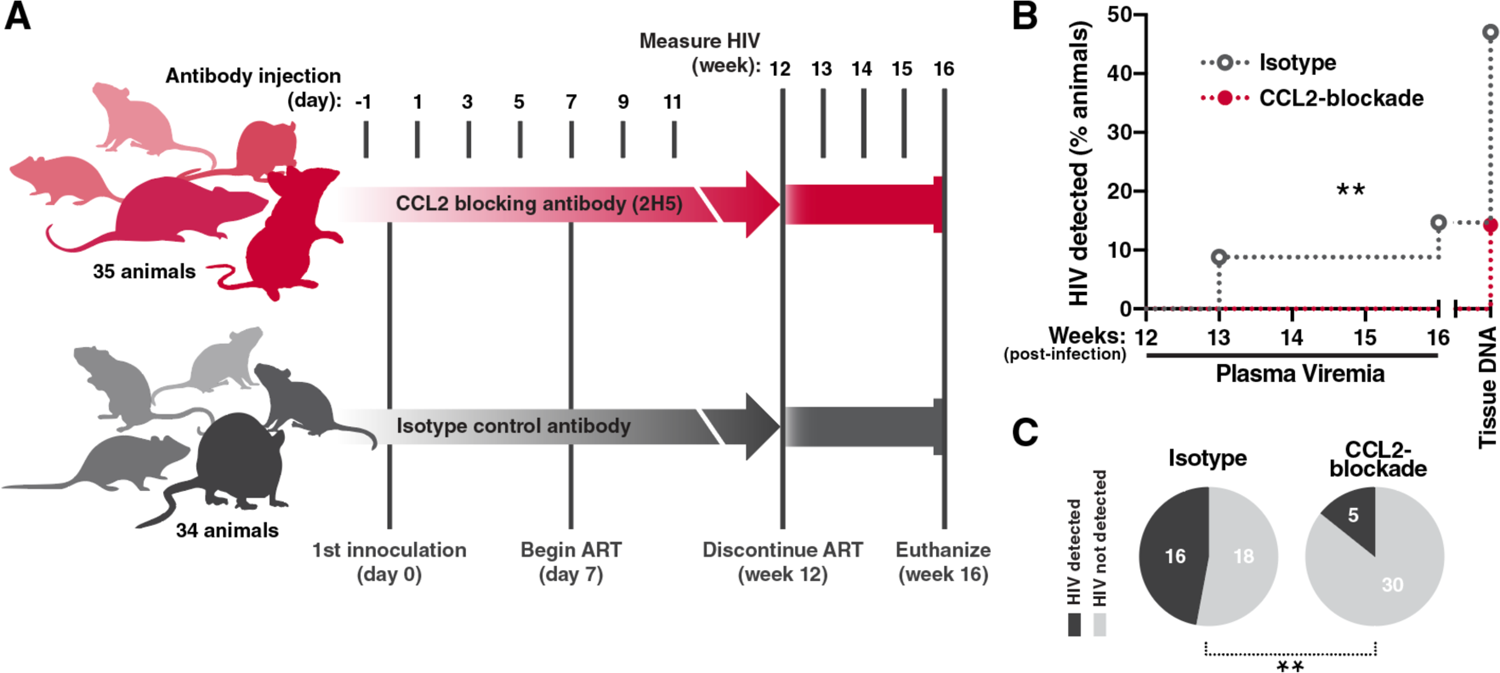
Blockade of CCL2 reduces seeding of the HIV reservoir in humanized mice. (**A**) Humanized mice were injected with 200 µg CCL2-blocking antibody (2H5) or isotype control beginning at 24 hours pre-infection, followed by 100 µg re-administered every 48 hours for a total of 12 days. Mice were challenged with HIV intrarectally at day 0 every 24 hours for 5 days and at day 7 were administered ART orally (chow infused). ART was maintained for 11 weeks and discontinued at week 12. Mice were bled weekly following ART-interruption and euthanized at week 16 post-infection. (**B**) HIV plasma RNA was quantified following ART-interruption and HIV DNA was measured in the spleen following euthanasia. CCL2-blockade decreased the detection of HIV (Log-rank test = 0.002) and (**C**) overall frequency of animals with detectable HIV (Fisher exact = 0.0041). Cumulative data shown from two independent experiments.

### CCR2/5+ cells harbor HIV provirus *in vivo*

To confirm that CCR2/5+ cells played a role in formation of the latent reservoir in people living with HIV on ART, we examined if CCR2/5+ CD4 T cells from such individuals were enriched in HIV proviruses. Fresh leukapheresis samples were obtained from nine ART-suppressed donors from the SCOPE (“Observational Study of the Consequences of the Protease Inhibitor Era”) cohort at the University of California San Francisco and San Francisco General Hospital (ClinicalTrials.gov Identifier: NCT00187512) (Figure 6––table supplement 2). From these samples, CD4 T cells were sorted into naïve (CD45RO-) and memory (CD45RO+) cell populations. The memory population was further separated based on the surface expression of CCR2 and CCR5 (Figure 6A).

**Figure 6:**
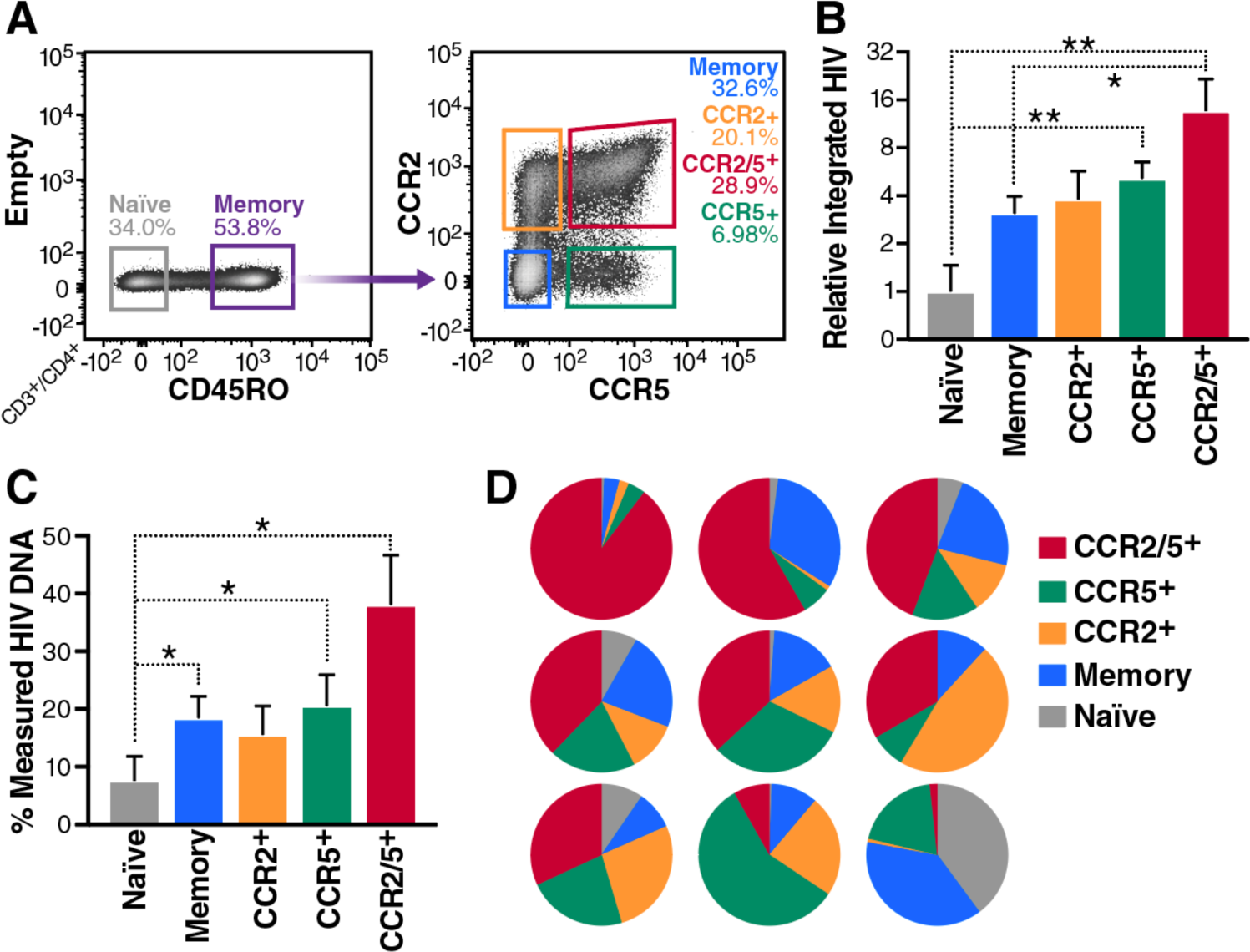
Integrated HIV proviral DNA is enriched in CCR2/5+ cells. Naïve, Memory (CCR-), CCR2+, CCR5+, or CCR2/5+ CD4 T cells were purified by cell sorting from leukapheresis samples obtained from ART-suppressed, HIV-infected donors. (**A**) Example of sorting gates from donor cells, pre-gated on live, single CD3+, CD4+ cells. (**B**) Amount of integrated HIV in sorted cell populations, normalized to mitochondrial DNA and expressed as fold increase over the amount detected in naïve cells. Cells were sorted as in panel A, genomic DNA was isolated, and provirus was measured using ALU-gag droplet digital PCR. (**C** and **D**) Distribution of HIV provirus among various cell populations from nine ART-suppressed HIV-infected donors. Results are shown as a percentage of total HIV detected from all subsets for each individual (n=9). Results are grouped by cell type in panel C and separated by individual in panel D. Error bars denote SEM, Friedman ANOVA with Dunn’s test for multiple comparisons, * p < 0.05, ** p < 0.01.

To quantify proviral levels, genomic DNA from sorted cells was isolated and subjected to ALU-Gag droplet digital qPCR to detect integrated HIV provirus. This technique is based on a nested PCR, in which an Alu-Gag amplicon is pre-amplified and a second internal set of primers + probe is used to detect HIV. Using this assay, we observed that CCR2/5+ cells contained significantly higher levels of integrated HIV than CCR2/5-memory cells or naïve cells (Figure 6 B-D). Additionally, by comparing the levels of HIV DNA in the sorted populations to that in the total population of sorted CD4 T cells, we were able to quantify the percentages of total HIV provirus in each cell population for each donor (Figure 6 C and D). Considerable individual to individual differences were observed. One individual exhibited a reservoir that was nearly 90% within the CCR2/5+ population, while in others this percentage was lower. The median percentage of HIV provirus within the CCR2/5+ cells was 37% (Fig 6D). Importantly, the CCR2/5+ cells had the highest proportion of integrated HIV of all the sorted subsets. These results suggested that CCR2/5+ cells formed a significant fraction of the latent HIV reservoir *in vivo*.

## DISCUSSION

Chemokines and chemokine receptors have long been recognized to play important roles in HIV biology. For example, the CCR5 and CXCR4 chemokine receptors function as coreceptors for HIV entry, ^46, 47^ while the beta-chemokines, MIP-1*α*, MIP-1*β*, and RANTES, act as potent inhibitors of R5-tropic virus infection due to their ability to bind to and downregulate membrane expression of CCR5 ^48^.

Various HIV proteins including HIV gp120, Tat, Nef, and p17 can activate expression of different chemokines including CCL2, CCL3, CCL4, and CCL5 involving effects on both infected and bystander cells ^14^. During HIV infection, CCL2 is prominently and rapidly expressed. However, CCL2 production is not protective for the host, rather this chemokine promotes HIV infection and spread both through the recruitment of new target cells (monocytes and CD4 T cells) and by enhancing HIV replication via CCR2 signaling. CCL2 plays an important role in the development of HIV-associated Neurocognitive Disorder (HAND) by enhancing blood-brain barrier permeability and recruiting macrophages and activated T cells resulting in a chronic inflammatory encephalitis^16–20^. Similarly, CCL2 has been linked to other chronic inflammatory responses occurring in multiple sclerosis, rheumatoid arthritis and various kidney diseases ^31, 49^.

Regarding HIV latency, the initial latent reservoir is formed within a few hours to a few days after HIV infection. Central memory CD4 T cells are recognized as an important, self-renewing subset of cells harboring latent provirus. The mechanisms underlying rapid seeding of the HIV reservoir in these long-lived memory CD4 T cells remain unclear. Our studies now suggest that high level production of CCL2 in tissue zones of HIV infection and inflammation recruits CCR2/5+ central memory CD4 T cells likely contributing to rapid establishment of the latent HIV reservoir.

### An additional mechanism of CCL2 induction by HIV

Cell-to-cell infection is fundamental to the pathogenesis and spread of HIV infection ^21–23, 50, 51^. Early in our studies of tonsillar lymphocytes, we noted similar patterns of antiviral drug inhibition for the production of CCL2 and type I interferons and the pyroptotic response that accompanies abortive HIV infection of nonpermissive bystander cells (inhibition by entry inhibitors and NNRTI’s but no inhibition by NRTI’s or integrase inhibitors). Pyroptosis and IFN1 production both involve IFI16 sensing of incomplete HIV reverse transcripts accumulating in abortively infected, nonpermissive bystander CD4 T cells present in lymphoid tissue ^25, 26^. These responses do not occur with blood CD4 T cells although coculture of blood and lymphoid tissue cells confers pyroptotic sensitivity to the CD4 T cells. ^21, 25, 26^. Like pyroptotic cell death, CCL2 production occurs in lymphoid, but not blood, CD4 T cells in our culture system (Figure 1 and Figure 1––figure supplement 1). Others have previously shown that IFI16 signaling via STING drives IRF3 phosphorylation and Type I IFN production ^27^; and further, that STING stimulates CCL2 production by activating STAT6 ^28^. When sgRNAs targeting IFI16 and STING were deployed with Cas9 RNPs to knockout these genes in lymphoid CD4 T cells, a partial decline in CCL2 production following HIV infection was observed. In contrast to the STING and IFI16 gene deletions, knockout of cGAS, a primary activator of STING ^29^, only trended to decreasing CCL2 production not reaching statistical significance.

Of note, the IFI16 and STING knockouts only partially ablate production of CCL2 in the HIV-infected CD4 T cells. This likely is due to the fact that CCL2 production during HIV infection occurs through multiple mechanisms. As noted above, HIV gp120 ^52–54^, Tat ^55–57^, Nef ^58^, and p17 ^59^ have all been shown to activate CCL2. Our findings now add an additional mechanism underlying CCL2 release during HIV infection involving IFI16 sensing of abortive reverse transcription products coupled with STING signaling.

### CCR2/5+ lymphocytes are primarily central memory cells in lymphoid tissue

CCL2 attracts cells that express CCR2 to areas of tissue inflammation. These cells can include monocytes, CD4 T cells, and NK cells. Of note, CCR2 and CCR5 share 73% sequence homology, are both found on the long arm of chromosome 17, and likely arose as a result of gene duplication ^60, 61^. While the circulating CCR2/5+ CD4 T cells in blood are predominantly effector memory cells ^32^, the CCR2/5+ cells in the lymphoid tissue display a central memory phenotype (Figs. 2 and 3). This pattern of chemokine receptor expression is intriguing in the context of HIV infection as it provides a potential pathogenic link between target cell recruitment (CCR2), infectability (CCR5), and rapid establishment of a long-lived latent reservoir within central memory T cells (CCR7).

### CCR2/5+ cells are targets of HIV infection

We find that sorted CCR2/5+ cells from unstimulated tonsils are permissive to HIV infection (Figure 4). However, it is important to sort these cells prior to infection before R5-tropic viruses induce a downregulation of surface CCR2 changing the cell’s phenotype (Figure 4D). We suspect that this loss of CCR2 expression is due to viral envelope interactions with CCR2. Non-infected bystanders (GFP-) as well as infected cells (GFP+) exhibit downregulation of CCR2 following exposure to R5-tropic, but not X4-tropic NL4-3. In addition to the homology shared by CCR2 and CCR5, previous studies have described CCR2 binding by HIV Env, and multiple HIV entry inhibitors block both CCR5 and CCR2—including TAK-779 and cenicriviroc ^62^. Thus, our observation of HIV envelope-mediated downregulation of CCR2 is likely explained by the similarity of CCR2 and the canonical CCR5 HIV co-receptor with envelope binding sufficing to trigger internalization of both receptors.

By attracting target cells to become productively infected, the CCL2/CCR2 axis could contribute to acute HIV pathogenesis amplifying productive, spreading infection. Additionally, CCR2/5+ cells may undergo latent infection facilitating early formation of the reservoir. Using an *in vitro* model to measure HIV reactivation, we find that CCR2/5+ cells harbor significant levels of latent provirus that can be induced by subsequent T cell receptor stimulation (Figure 4E).

### Mucosal origins of CCR2/5+ cells

As the primary route of HIV infection is through mucosal tissues, the integrin expression profile of the CCR2/5+ population is particularly interesting. These cells express high surface levels of the CD49d, CD29, and α4β7 integrins (Figure 3) that play a key role in mucosal trafficking ^63^. CCR2+ CD4 T cells are also recruited to the ileum in inflammatory bowel disease ^64^. Thus, CCR2/5+ CD4 T cells are likely mucosal in origin, or alternatively home to mucosa or mucosa-associated lymphoid tissue (MALT). These cells may be involved in establishment of reservoirs in MALT even before migration into secondary lymphoid organs.

### In Vivo Blockade of CCL2 reduces HIV reservoir formation

We next sought to test the biological relevance of CCL2 action *in vivo* in the formation of the latent HIV reservoir in BLT-humanized mice ^44, 45, 65^ and our studies indicated that antibody-mediated blockade of CCL2 significantly reduced the formation of the HIV reservoir, as measured by decreased detection of plasma viremia following ART withdrawal and total detectable HIV at necropsy (Figure 5). CCL2 blockade was administered one day prior to HIV exposure. We also intentionally selected a short infection period of only 6 days before initiation of ART to evaluate the establishment of the most rapidly formed stable reservoir.

Our planned follow-on investigations in the BLT-humanized mouse model including lengthening the time of infection could not be performed due to the Presidential Executive Order banning government scientists (including K.J.H.) from using human fetal tissue in research. Despite these limitations, our findings are consistent with anti-CCL2 blockade decreasing early seeding of the HIV reservoir *in vivo* and merit further examination.

### CCR2/5+ cells serve as hosts for a substantial fraction of the latent HIV reservoir in humans

Long-term effective ART contracts the population of HIV-infected cells around a core reservoir of cells infected prior to treatment. Our findings indicate that on average CCR2/5+ cells harbor ∼35% of the reservoir when blood cells are analyzed from HIV-infected individuals on ART. However, we also noted marked differences in the fraction of the reservoir residing in CCR2/5+ cells among the 9 individuals analyzed (Fig 6D). In one individual, nearly 90% of the latent reservoir was found in the CCR2/5+ cells. These findings suggest that CCL2 may play an important role in latent reservoir seeding in many but not all infected individuals. It is certainly possible that the underlying mechanism of infection and level of inflammation in lymphoid tissues may vary—impacting levels of CCL2 that are released, and the extent to which CCR2/5+ chemoattraction occurs. Additionally, the duration of infection prior to treatment may allow for spread of the reservoir, and although CCR2+ cells may be initially favored, the phenotype of the reservoir likely broadens during chronic infection. CCR2-negative cells also harbored HIV provirus in all assayed individuals (Figure 6). Therefore, CCR2 expression certainly does not perfectly mark the HIV reservoir *in vivo*.

A limitation of most reservoir studies, including this one, is that human samples are often limited to peripheral blood donated by HIV-infected individuals, and therefore fail to sample the tissue reservoir. However, circulating cells of the blood originate from tissue sites. The effector memory CCR2/5+ cells we find containing HIV provirus are likely daughters of the initially infected lymphoid tissue-derived CCR2/5+ central memory cells.

Together, our results demonstrate that lymphoid tissue CD4 T cells rapidly produce CCL2 as part of an innate inflammatory response launched during HIV infection. Among multiple mechanisms, IFI16 sensing of HIV DNA and signaling via STING contributes to rapid release of the CCL2 chemokine that can quickly recruit CCR2/5+ central memory cells to the zone of HIV-associated inflammation. Within these infection zones, some cells may become productively infected while others undergo latent infection and yet others die. Finally, CCR2/5+ central memory cells take up residence in peripheral lymphoid organs, harboring latent HIV provirus, contributing to the persistence of the stable reservoir. Our findings argue for a role of the CCL2/CCR2 axis in early seeding of the latent reservoir illustrating how the innate immune response may, in this case, promote viral persistence.

## MATERIALS AND METHODS

### Experimental Model and Subject Details Human samples

Blood from HIV-infected individuals was obtained from volunteers participating in the SCOPE cohort ^66^. Participants gave their informed consents as part of the SCOPE cohort. Specific characteristics of these participants and their ART regimens are summarized in Table S2.

### Humanized TKO-BLT mice

Male and female C57BL/6 Rag2-/-γc-/-CD47-/-(TKO) mice were humanized using the bone marrow, liver, thymus (BLT) method as previously described ^44^. All animal studies were performed after approval of the animal study protocol by the AAALAC-accredited Rocky Mountain Laboratories, National Institute of Allergy and Infectious Diseases, National Institutes of Health (USA) Institutional Animal Care and Use Committee. The mice were housed under specific pathogen-free conditions, properly anesthetized during procedures and monitored daily. Donor tissues for humanization were obtained with informed consent following the guidelines and regulations of NIH and the Office of Human Subjects Research Protection. All humanized mouse studies were initiated prior to the Presidential executive order banning government research using fetal tissue.

Humanized mice were treated intraperitoneally with 200 µg anti-CCL2 antibody (2H5) or isotype control 24 hours prior to infection followed by 100 µg administered every 48 hours for a total of 12 days. HIV-1 JR_CSF_ stocks were prepared and inoculated as previously described (55). The mice were challenged with 5×10^4^ TCIU of HIV via the anal route at day 0 and repeated every 24 hours for 4 additional days. At day 6 post-infection, the mice were bled retroorbitally. At day 7, the mice were free-fed dyed red ART-infused chow (Modified LabDiet® PicoLab® Mouse Diet 20, 5058; supplemented with 1250 ppm Emtriva (ematricitabine), 1630 ppm Viread (tenofovir disproxil-fumerate), and 10,688 ppm Isentress (raltegravir)) until 12 weeks post infection, when ART was discontinued. Mice were bled at 6 weeks post-infection to test for suppression of viremia and weekly following ART-interruption until the euthanasia time-point at 16 weeks post-infection.

### Primary-cell cultures

Specimens derived from HIV-negative human blood and tissue were de-identified before receipt by the laboratory and are thus exempt from human subject research per the UCSF Human Research Protection Program Institutional Review Board. Human healthy tonsils and spleens were obtained from the Cooperative Human Tissue Network (https://www.chtn.org). HIV-negative donor blood was obtained from Vitalant (https://www.vitalant.org).

Human lymphoid aggregate culture (HLAC) prepared from tonsil or spleen was cultured in HLAC medium: RPMI supplemented with 15% heat-inactivated fetal bovine serum (FBS), 100 mg/ml gentamicin, 200 mg/ml ampicillin, 1 mM sodium pyruvate, 1% nonessential amino acids, 2 mM L-glutamine, and 1% fungizone at 37°C in 5% CO_2_ incubator.

Concentrated white blood cell preparations from healthy volunteers were obtained from Vitalant. PBMCs were cultured in RPMI supplemented with 10% FBS, 1000 U/ml Penicillin and 1 mg/ml Streptomycin and 2 mM L-glutamine, at 37°C in 5% CO_2_ incubator.

### Cell line

HEK293T cells were transfected with various molecular clones of HIV to produce high titer virus preparations. Cells were cultured in DMEM supplemented with 10% FBS, 1000 U/ml Penicillin and 1 mg/ml Streptomycin and 2 mM L-glutamine, at 37°C in 5% CO_2_ incubator.

### Virus strains

HIV molecular clones (pNL4-3-GFP, pBaL-GFP, and pNL4-3-Luciferase) were purified from E. coli and used to transfect HEK293T cells.

### Primary cell isolation & culture

Fresh human tonsil tissue or splenic tissue was processed and cultured as previously described^24^. Briefly, single-cell suspensions were prepared by mechanical disruption of the tonsil or spleen, followed by sequential passage of cells through 70 µm then 40 µm strains. Single-cell suspensions were purified by density gradient centrifugation using Ficoll-Hypaque (GE), the mononuclear cell layer was isolated and washed twice with FACS buffer and then analyzed by flow or mass cytometry, or alternatively cultured in HLAC medium. PBMCs were similarly purified by density gradient centrifugation using Ficoll-Hypaque.

### CD4 T cell isolation

CD4 T cells were purified from lymphoid tissue or PBMC-derived mononuclear cells by negative magnetic depletion using the EasySep Human CD4+ T Cell Enrichment Kit (Stem Cell), per the manufacturer’s protocol.

### HIV infection (overlay model)

Overlay infections were carried out as previously described ^21^. Briefly, HEK293T cells were seeded in 24-well tissue culture plates at a density of 1.6 x 10^5^ cells/well and transfected with 50-100 ng/well of pNL4-3-GFP using Fugene HD (Promega) transfection reagent, per the manufacturer’s protocol. The medium was replaced after 16 hours, and CD4 T cells, purified as described above, were added. Cells or supernatants were harvested at time points indicated for each experiment.

### HIV infection (spinoculation model)

Concentrated HIV was produced by transfecting 5-10 x 10^6^ HEK293T cells per T150 flask with pNL4-3.GFP, pBAL.GFP, or pNL4-3.Luciferase, using Fugene HD per the manufacturer’s protocol. The medium was replaced at 16 hours after transfection and supernatant harvested at 48- and 72-hours post-transfection. Supernatant containing viruses was passed through a 0.22 µm filter and virus was pelleted at 25,000 x g for 2 hours at 4°C. Following resuspension of the pellet in RPMI, the concentrated viruses were aliquoted and frozen at −80 °C. Virus concentration was determined by quantitation of p24^gag^ as previously described ^21^. 100 ng of virus was generally added to the cell populations cultured at 1 x 10^6^ cells/mL in 100 µL in 96-well round-bottom tissue culture plates. The plate was spun at 900 x g for 2 hours at room temperature and then cultured at 37°C in 5% CO_2_ incubator.

### Analysis of cytokine expression

Quantitative RT-PCR was used to assess cytokine mRNA expression. Cells were harvested 18-48 hours following culture or infection as described for the various experiments, and RNA was purified using RNAeasy Kit (Qiagen) per the manufacturer’s protocol with inclusion of DNAse treatment. Purified RNA was quantified, and cDNA was prepared using the SuperScript® III First-Strand Synthesis kit (Thermo Fisher). RNA, oligo(dT)_20_ and dNTPs were combined and incubated at 65°C for 5 minutes, then cooled to 4°C. RT buffer, MgCl_2_, DTT, RNaseOUT™, and SuperScript® III RT were added per the manufacturer’s protocol and incubated at 50°C for 50 minutes. The reaction was terminated by heating to 85°C for 5 minutes and cDNA was stored at −20°C. Following cDNA production, quantitative PCR was carried out using Taqman Gene Expression Assays (Thermo Fisher, see Key Reagents Table for specific Assay IDs) per the manufacturer’s protocol. Both GAPDH and 18S mRNA were measured, and the geometric means of these housekeeping genes were used to normalize expression (delta cycle threshold) of query transcripts.

Secreted CCL2 protein was measured using the Meso Scale Discovery (MSD) platform. Supernatants from cells cultured as described were harvested and clarified by centrifugation at 1000 x g for 10 minutes. Undiluted supernatant was applied to the V-PLEX Human MCP-1 Kit (Meso Scale Discovery) and CCL2 was measured using a recombinant standard, per the manufacturer’s protocol.

### Cas9 RNP genomic editing

Nucleofection was performed using *in vitro* assembled Cas9RNP nucleofection as previously described ^67^. Recombinant Cas9 protein derived from *S. pyrogenes* engineered with two nuclear localization signals and an HA tag on the C-terminus was obtained from the QB3 Macrolab at the University of California Berkeley. Multiple guide RNAs were tested, and two were selected for each target gene that showed high knockdown efficiency in multiple donors (see Key Resources Table for guide sequences). Guide RNA and tracrRNA suspended in 10mM Tris-HCL (pH 7.4) were mixed at 1:1 ratio, at a final concentration of 40 µM and incubated for 30 minutes at 37°C. The guide RNA mixture was added to 40 µM recombinant Cas9 and incubated at 37°C for 15 minutes to allow for assembly of the guide RNA:Cas9 RNPs.

HLAC cells were stimulated for 3 days with plates coated with 10 µg/mL anti-CD3 (UCHT1, Tonbo Biosciences) and 10 µg/mL anti-CD28 (CD28.2, Tonbo Biosciences). CD4 T cells were purified as described above and 3 x 10^5^ CD4 T cells were pelleted in a V-bottom plate and resuspended in 20 µL P3 buffer, from the P3 Primary Cell 96-well Nuclofector kit (Lonza). 3 µL of the assembled Cas9 RNPs were added to the cells, and the mixture was transferred to a 96-well reaction cuvette.

The cuvettes were loaded into the 4D-Nucleofector (Lonza) and electroporated using program EH-115. 80 µL of pre-warmed HLAC medium was gently added to each well and the cells were allowed to recover for 30 minutes at 37°C, before re-stimulating overnight on plates as above. Following Cas9RNP-mediated editing, the cells were removed from stimulation and rested for one week before infection or subsequent analyses. Knockout efficiencies were measured using immunoblotting or intracellular flow cytometry.

### Immunoblotting

Cell lysates were prepared by addition of Laemmli Lysis Buffer and heat denaturization at 90°C for 30 minutes. Lysates were electrophoresed on a 4-12% Bis/Tris gel (Thermo Fisher) at 120V for 90 minutes. Proteins were transferred to a PVDF membrane using an iBlot 2 apparatus per the manufacturer’s protocol (Default program P1). Following transfer, the membrane was blocked with 5% w/v dry skim milk dissolved in PBS + 0.1% Tween-20 (PBS-T) for 1 hour at room temperature with rocking. Blocked membranes were incubated with primary antibodies (see Key Resources Table), diluted in blocking buffer (1:1,000) and incubated with rocking overnight at 4°C. The following day, membranes were washed in PBS-T and incubated with HRP-conjugated secondary antibody diluted 1:10,000 in blocking buffer for 2 hours at room temperature with continuous rocking. After secondary incubation, the membranes were washed and TMB substrate was added. The reaction was allowed to proceed for 5 minutes, and light was measured by exposure to film.

### Flow cytometric analysis & cell sorting

Flow cytometry was performed at the Gladstone Institutes Flow Cytometry Core Facility using two instruments: an LSR II (BD Biosciences, San Diego, CA, US) and FACSAria II (BD Biosciences, San Diego, CA, US). Cells were resuspended in PBS and Zombie Live/Dead (Biolegend) stain was added per the manufacturer’s protocol. Subsequently, FACS buffer (PBS + 2mM EDTA + 5% BSA) was added as well as antibodies for cell surface staining (see Key Resources Table). Cells were washed and sorted live; or fixed in 2% paraformaldehyde (PFA) prior to analysis. For intracellular antigen detection, following cell surface staining, cells were permeabilized using the Foxp3/Transcription Factor Staining Buffer Set (Thermo Fisher) following the manufacturer’s protocol.

Fluorescence-activated cell sorting was performed on unfixed live cells, following isolation and surface staining as described above. Single cells from a lymphocyte scatter gate were gated as “live” (zombie low/negative, see Key Resources Table), and subsequently as CD3+/4+ T cells. CD4+ T cells were then sorted as naïve (CD45RO-) or memory (CD45RO+) and further sub-divided as CCR2-/5-, CCR2+/5-, CCR2-/5+, or CCR 2+/5+, (see Figure 6 for example). Cells were sorted into ice cold HLAC medium for functional studies as described in the text; or sorted into ice cold PBS, pelleted, and genomic DNA isolated for HIV provirus measurements.

### Mass cytometric analysis

Mass cytometric analyses was performed as previously described ^68^. Cells were isolated from fresh PBMCs or HLAC as described above and stained with a panel of lanthanide metal-conjugated antibodies (See Table S1). Antibody staining was performed in a volume of 100 μL for 45 min, treated with 139In-DOTA-maleimide (Macrocyclics) to label dead cells, and fixed overnight with PBS + 2% PFA. Cells were permeabilized with FoxP3 Fix/perm buffer (ThermoFisher) per the manufacturer’s protocol, then labelled with 1:4,000 191/193Ir DNA intercalator (Fluidigm). Cells were washed, resuspended in distilled water, and analyzed on a CyTOF2 (Fluidigm) at the UCSF Parnassus Flow Cytometry Core. EQ calibration beads (Fluidigm) were included and used for normalization across runs.

### Chemotaxis measurement

CCR2/5+ CD4+ T cells were purified by FACS and suspended at 1 × 10^6^ cells/ml in RPMI with 0.5% BSA and 25 mM HEPES. 100 μl of sorted cells were added to each transwell insert with 6.5-mm-diameter membranes containing 3.0-μm pores (Corning). The cells were pre-incubated for 30 min at 37°C, and the inserts transferred into wells with chemotaxis medium containing either CCL2 (1 μg/mL, PeproTech) or medium alone. Cells in the lower wells were harvested after 3 hours at 37°C, and the migrated cells were counted using AccuCount Particles on an LSR II cytometer.

### In vitro latency

As previously described, purified CD4 T cells were initially infected and then rested in the presence of ART to establish *in vitro* latency ^43^. Briefly, tonsil-derived purified resting (unstimulated) memory CD4 T cells were sorted based on their expression of CCR2 and 5 as described above. 100 ng of purified NL4-3-Luciferase was added to the sorted cells and they were spinoculated as described above and 5 µM saquinavir was added to limit infection to a single round. Following 5 days of culture the cells were stimulated with anti-CD3/CD28 beads per the manufacturer’s protocol, in the presence of 30 µM raltegravir. 24 hours after stimulation, luciferase activity was measured as previously described ^43^.

### HIV Alu-Gag droplet digital qPCR

Nested Alu-gag qPCR was performed, as previously described ^69, 70^ with minor modifications. Briefly, genomic DNA was isolated using DNeasy kits using RNAse-treatment step (Qiagen) per the manufacturer’s protocol. A primary PCR to amplify integrated provirus used a forward ALU-targeting primer at a concentration of 100 nM and HIV-1 gag reverse primer at 600 nM. Pre-amplification was performed with Taq polymerase (Qiagen) for 20 cycles. 1 µL of the above PCR product was combined with 10 µL Gene Expression Master Mix (Applied Biosciences), 8 µL water,1 µL probe mix (Pre-made 20X mix: 5µM MH531 & MH532 primers, 4µM LRT probe). The PCR mixture was partitioned using a QX200 droplet generator (Bio-Rad), and the droplets were amplified for (95° for 15s, 60° for 30s x 35 cycles). Positive and negative droplets were then detected on a QX200 droplet reader (Bio-Rad), and the results quantified using QuantaSoft software.

### Measurement of HIV in TKO-BLT humanized mice

Plasma HIV RNA was isolated with the QIAamp Viral RNA Kit (Qiagen) and quantified using the Abbott RealTime HIV-1 Amplification Reagent Kit following the manufacturer’s protocols. Depending on the available sample volume, the detection limit was 375 cp/mL during follow-up or 150 cp/mL at the time of euthanasia. For the quantification of HIV DNA, genomic DNA of splenocytes was isolated using QIAamp DNA blood mini kit (Qiagen) and subjected to a probe-based real-time PCR approach, as previously described ^71, 72^. Briefly, HIV-specific and control (hCD3) DNA sequences were pre-amplified (12 cycles) in a TProfessional TRIO Thermocycler (Biometra). The pre-PCR amplicons were subjected to quantitative real-time PCR analysis using the Rotor-Gene Probe PCR Kit (Qiagen) performed in a Rotor-Gene Q instrument (Qiagen). Dual labelled probes were used for CD3 (YAK-BHQ1) and HIV-DNA (6Fam-BHQ1) detection. Plasmids encoding the corresponding amplicon regions or gDNA of cells harboring HIV-LTR sequences were used as standard curves.^72^

### QUANTIFICATION AND STATISTICAL ANALYSIS

Statistical details of individual experiments, including number of independent donors, mean values, standard error of the mean (SEM), and p values derived from statistical tests are described in the figure legends and specified in the figures. Statistical analyses were performed using GraphPad Prism software versions 7 & 8. p values ≤ 0.05 were considered statistically significant. For two-way column analyses, a Student’s two-tailed *t*-test was used. ANOVA tests were used for multiple comparisons with post-tests for multiple comparisons (as noted within). Asterisk coding in figures is as follows: * p ≤ 0.05; ** p ≤ 0.01; *** p ≤ 0.001. Data are presented as means with error bars indicating SEM unless otherwise stated.

## DATA AND CODE AVAILABILITY

This study did not generate datasets.

## ACKNOWLEDGMENTS

The authors thank Jane Srivastava and Nandhini Raman of the Gladstone Flow Cytometry Core, the UCSF Diabetes Research Center for use of their CyTof instrument, the UCSF Parnassus Flow Cytometry Core for CyTOF assistance, John C.W. Carroll for graphic arts, Francoise Chanut for editorial assistance, and Robin Givens for administrative assistance.

## SUPPLEMENTAL FIGURES

**Figure Supplement 1:**
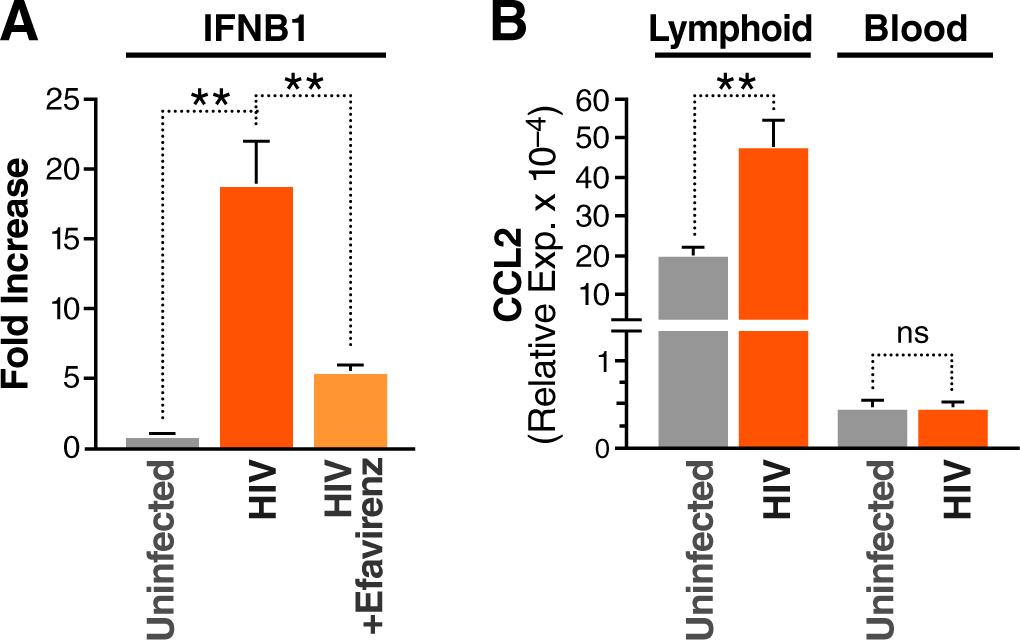
Cytokine responses to HIV by purified CD4 T cells. (**A**) Type 1 interferon is rapidly induced by lymphoid tissue CD4 T cells following HIV infection. Tonsillar CD4 T cells were purified by negative bead selection and infected with HIV by overlay infection for 18 hours ^21^. Production of IFNB1 RNA was measured by qPCR and plotted as fold increase relative to uninfected control. Treatment with efavirenz during overlay infection blocked induction of IFNB1. (**B**) CD4 T cells purified from PBMCs do not produce CCL2 in response to HIV. CD4 T cells were purified by negative bead selection from healthy donor PBMCs or tonsils and infected with HIV by overlay infection for 32 hours. Additionally, lymphoid tissue-derived CD4 T cells express substantially more CCL2 transcripts at baseline than PBMC T cells. Shown as level relative to housekeeping genes (geometric mean of GAPDH and 18S RNA). Experiments included 2-3 tonsil donors and were repeated at least twice. Error bars denote SEM (* p < 0.05, ** p < 0.01).

**Figure Supplement 2:**
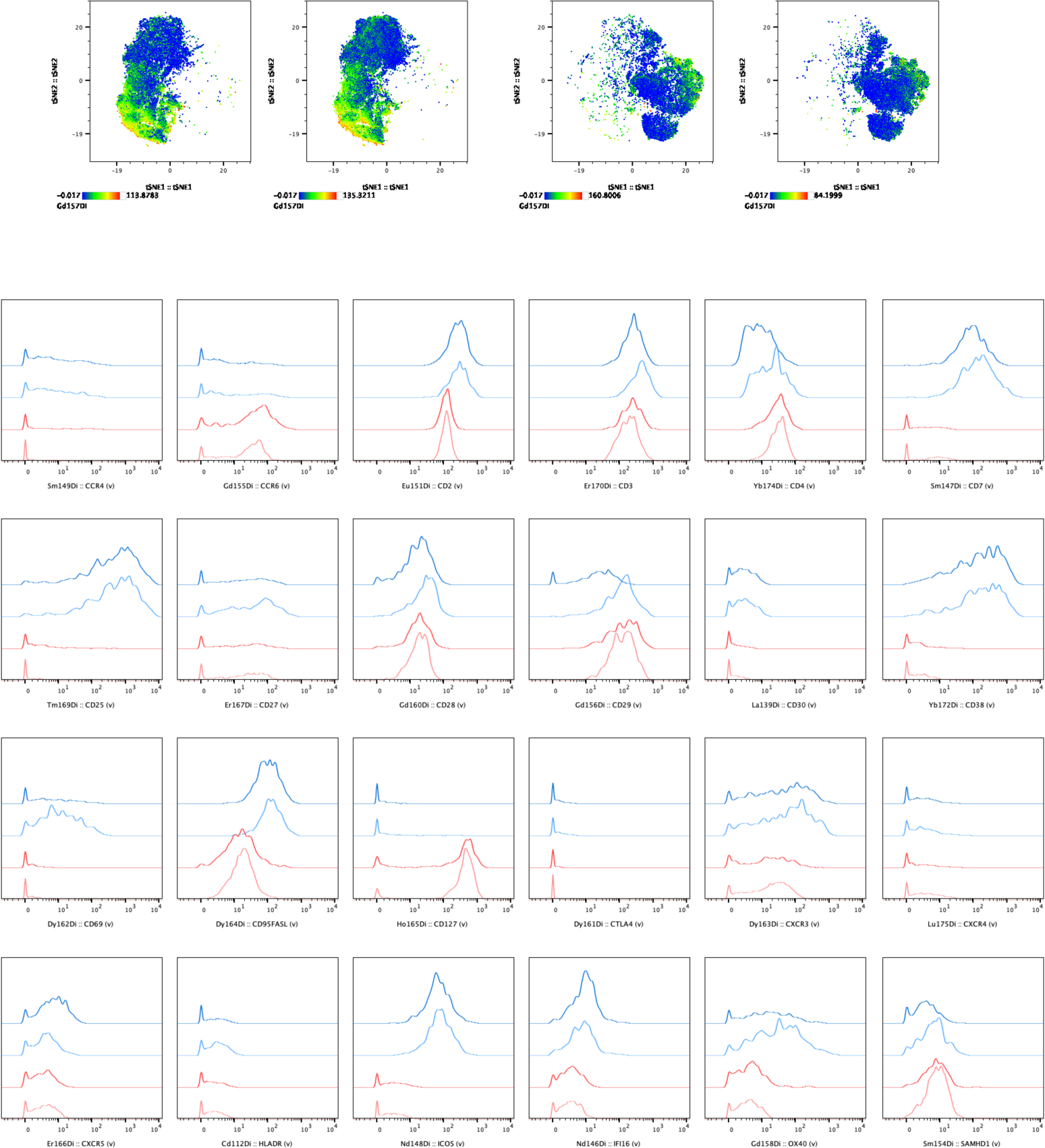
Comparison of blood and lymphoid tissue CCR2/5+ cells by mass cytometry. Fresh mononuclear cells were isolated from healthy donor PBMCs or tonsils and analyzed for expression of 38 markers by mass cytometry (n=10 donors of each tissue). (Top) tSNE dimensional reduction visualization was performed, colored by surface CCR2 expression. (Bottom) Cells were gated for CCR2/5 expression as in Figure 2. Shown here are a panel of markers which were different or similar between lymphoid tissue (blue) and blood-derived (red) CCR2/5+ cells.

**Figure Supplement 3:**
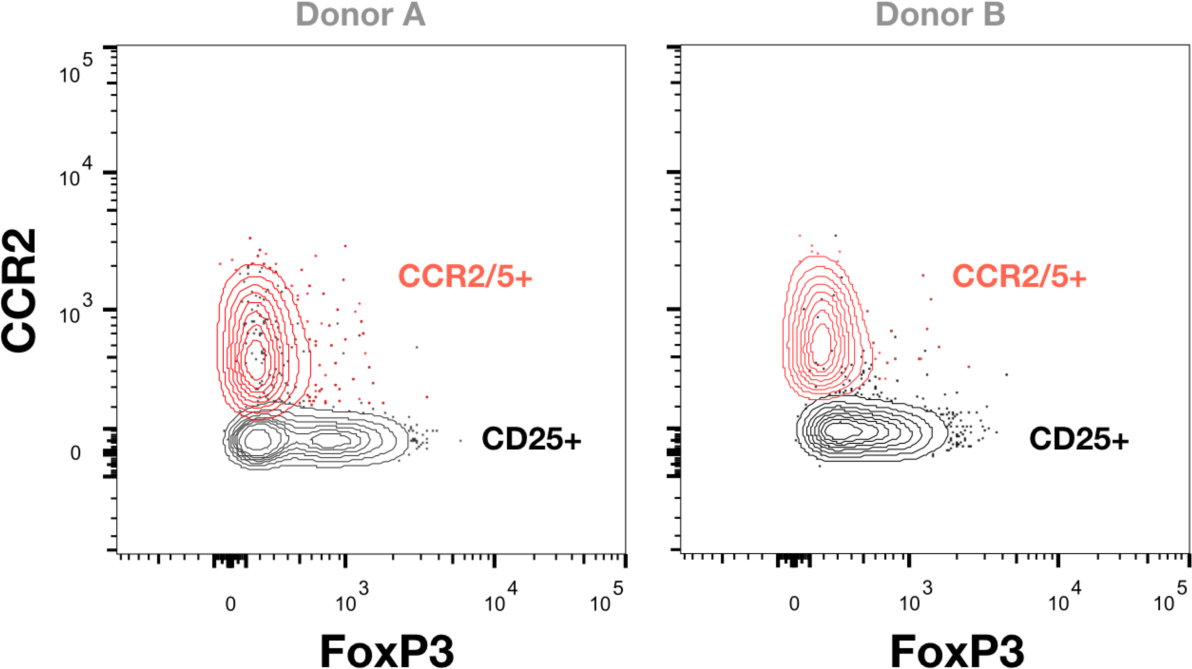
CCR2/5+ cells lack FoxP3 expression. CD4 T cells were purified by negative bead selection from two healthy donor PBMCs and surface stained, then fixed and permeabilized using FoxP3 Fix/Perm kit (eBiosciences) per the manufacturer’s protocol. Cells are gated on live singlets, CD3+, CD8-, CD4+, CD45RO+, and CD25+ (red) or CCR2/5+ (black).

**Figure Supplement 4:**
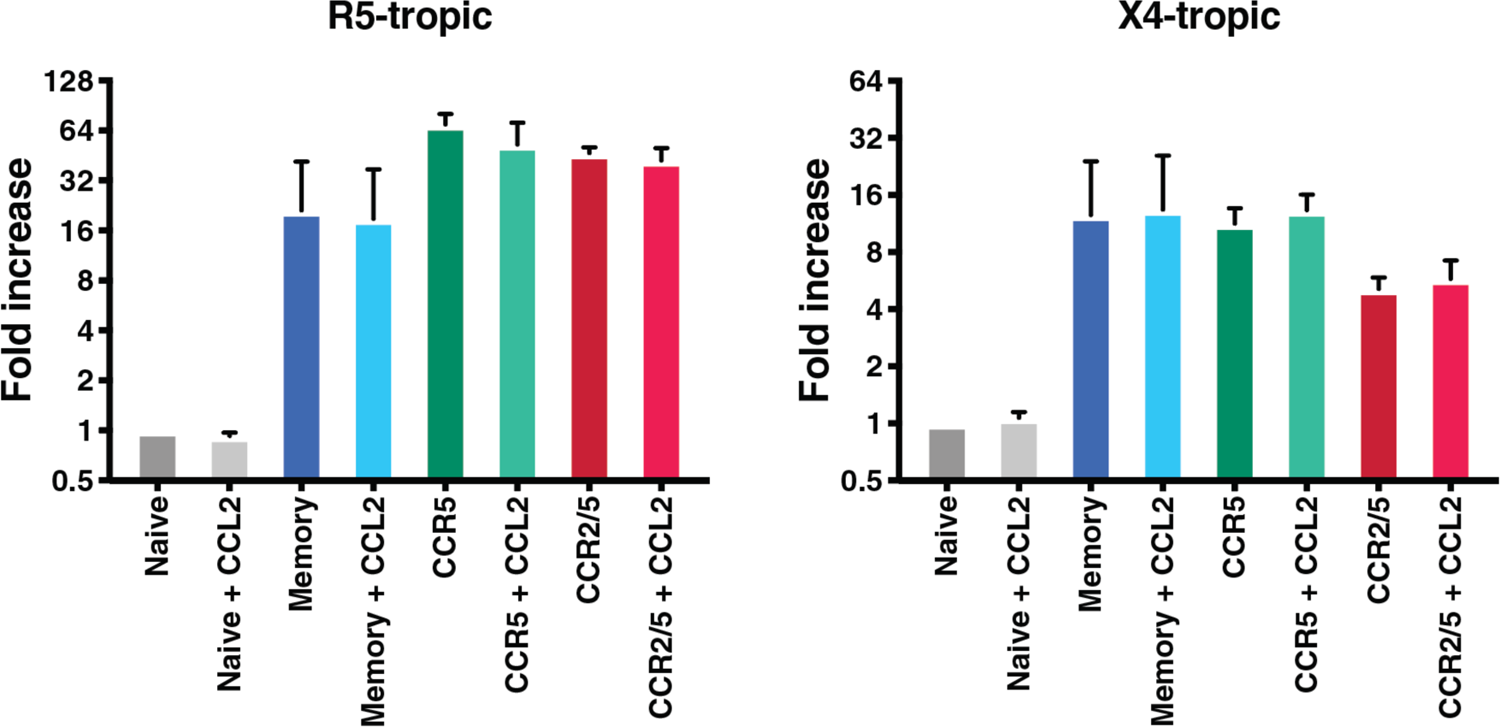
Addition of CCL2 does not enhance HIV infection in lymphoid tissue CD4 T cells. To test the effect of CCL2 signaling on HIV infection, cells were sorted from healthy tonsils as in Figure 4, and spinoculated with 100 ng GFP-reporter NL4-3 HIV-1 (X4-tropic) or BaL.NL4-3 HIV-1 (R5-tropic) followed by culture with or without CCL2 (5 µg/mL) (Peprotech). 48 hours later, levels of infection were measured by flow cytometric assessment of GFP+ cells and compared to infection levels in naïve cells. No significant differences in infection were observed in any of the cell populations treated with CCL2. Experiments included 2 tonsil donors and were repeated twice.

**Figure Supplement 5:**
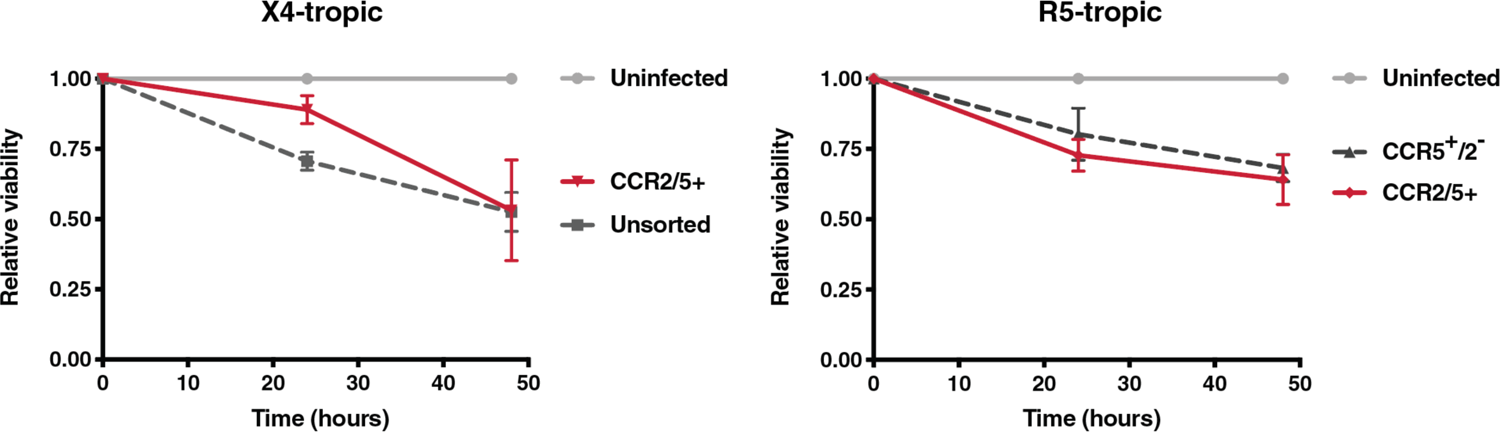
CCR2/5+ cells succumb to HIV infection at a similar rate as CCR5+ cells. To measure the susceptibility of CCR2/5+ cells to cell death following HIV infection, CD4 T cells were sorted and overlay infected with 50 ng NL4-3 HIV-1 (X4-tropic) or BaL.NL4-3 HIV-1 (R5-tropic), and live cells were quantified by normalizing to fluorescent beads at 24 and 48 hours post-infection. Experiments included 2 tonsil donors and were repeated twice.

**Table Supplement 1.**
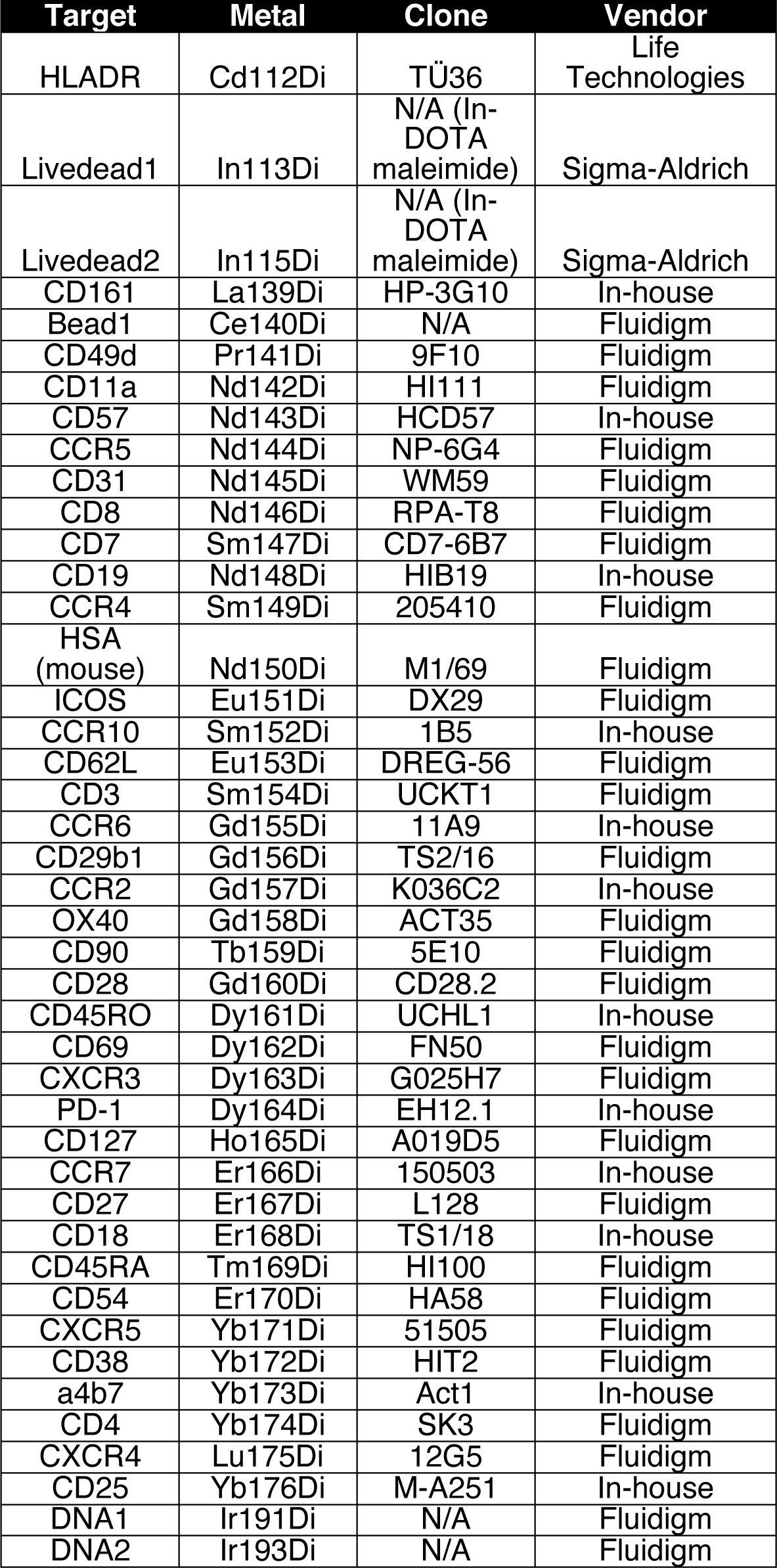
Mass Cytometry Panel

**Table Supplement 2:** **Characteristics of HIV-1 infected study participants** (Related to Figure 5 and STAR methods). Nine HIV-1-infected individuals, who met the criteria of suppressive ART (<50 copies/ml) for at least of six months, and a CD4+ T cell count of at least 500 cells/mm^3^ at time of study were enrolled. The participants were recruited from the SCOPE cohorts ^66^ 3TC, lamivudine; ABC, abacavir; ATV, atazanavir; COBI, cobicistat; DRV, darunavir; ETV, etravirine; FTC, emtricitabine; RGV, raltegravir; RPV, rilpivirine; RTV, ritonavir; TDF, tenofovir; TCV, tivicay.

| Participant ID | Age | Sex | Ethnicity | CD4 count (cells/ul) | ART regimen | ART start year |
| --- | --- | --- | --- | --- | --- | --- |
| 1695 | 39 | Female | Mixed | 556 | FTC/TDF, ATV, RTV | 2010 |
| 2021 | 51 | Male | Black | 540 | RPV/TAF/FTC | 1999 |
| 2146 | 71 | Male | Mixed | 204 | DRV/COBI, TCV, DES | 1995 |
| 2161 | 68 | Male | White | 687 | 3TC, RTV, DRV, TCV | 2005 |
| 2168 | 58 | Male | African American | 246 | ABC/TCV/3TC | 2000 |
| 2375 | 71 | Male | White | 727 | EFV/TDF/FTC | 1995 |
| 2402 | 40 | Male | African American | 765 | RTV, DRV, RPV/TDF/FTC | 2006 |
| 2461 | 63 | Male | White | 496 | TCV, RPV | 1999 |
| 2511 | 48 | Male | White | 334 | EFV/TDF/FTC, RGV | 2010 |

## KEY RESOURCES TABLE

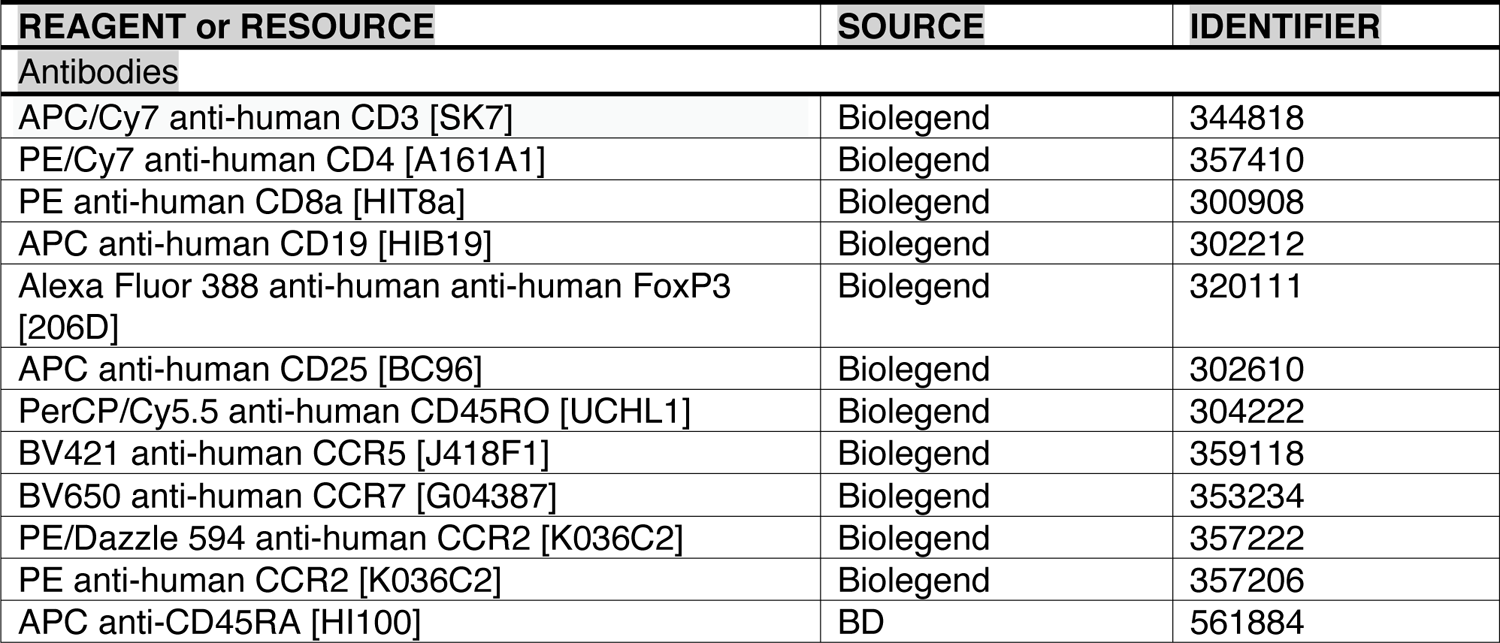
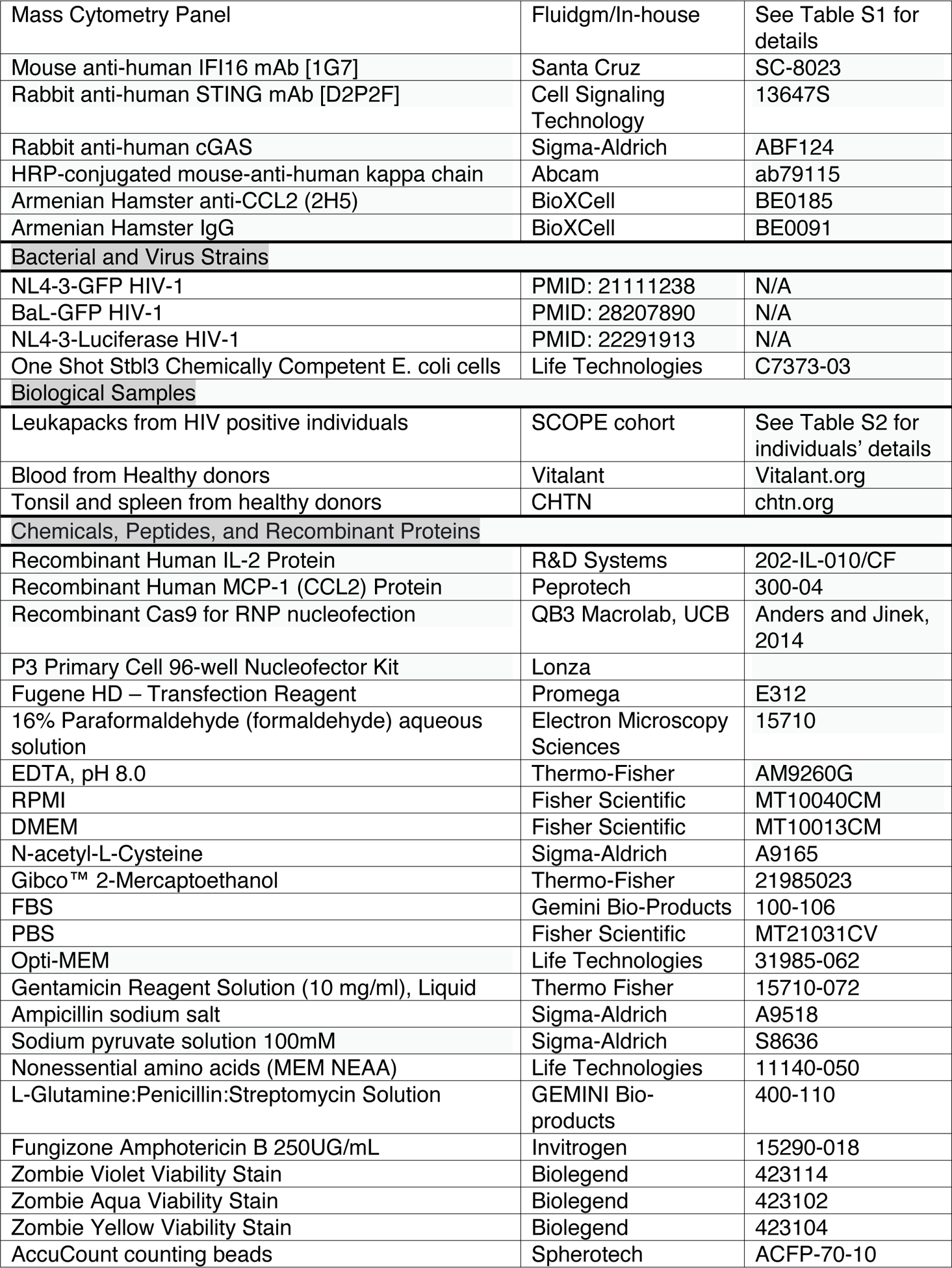
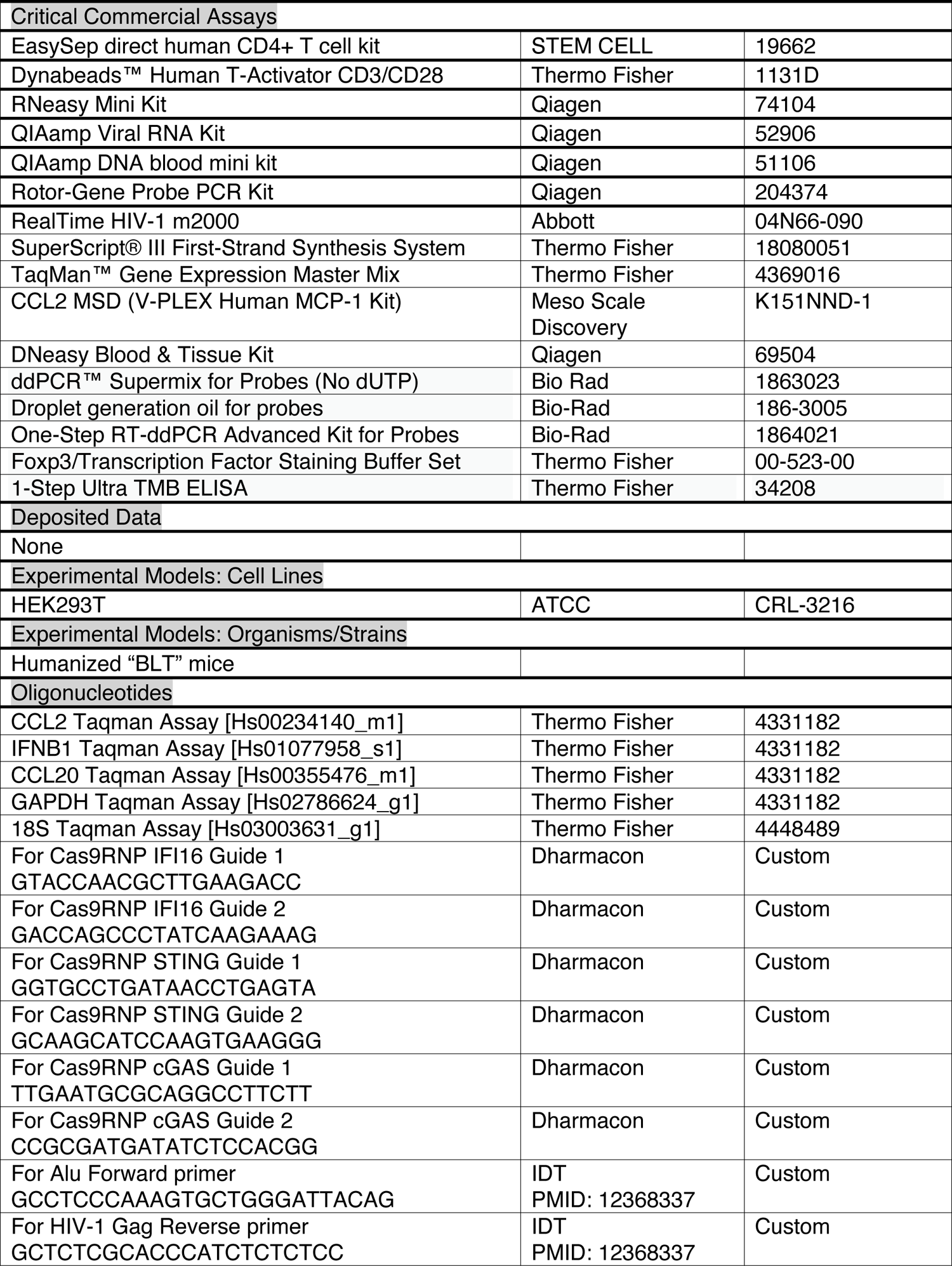
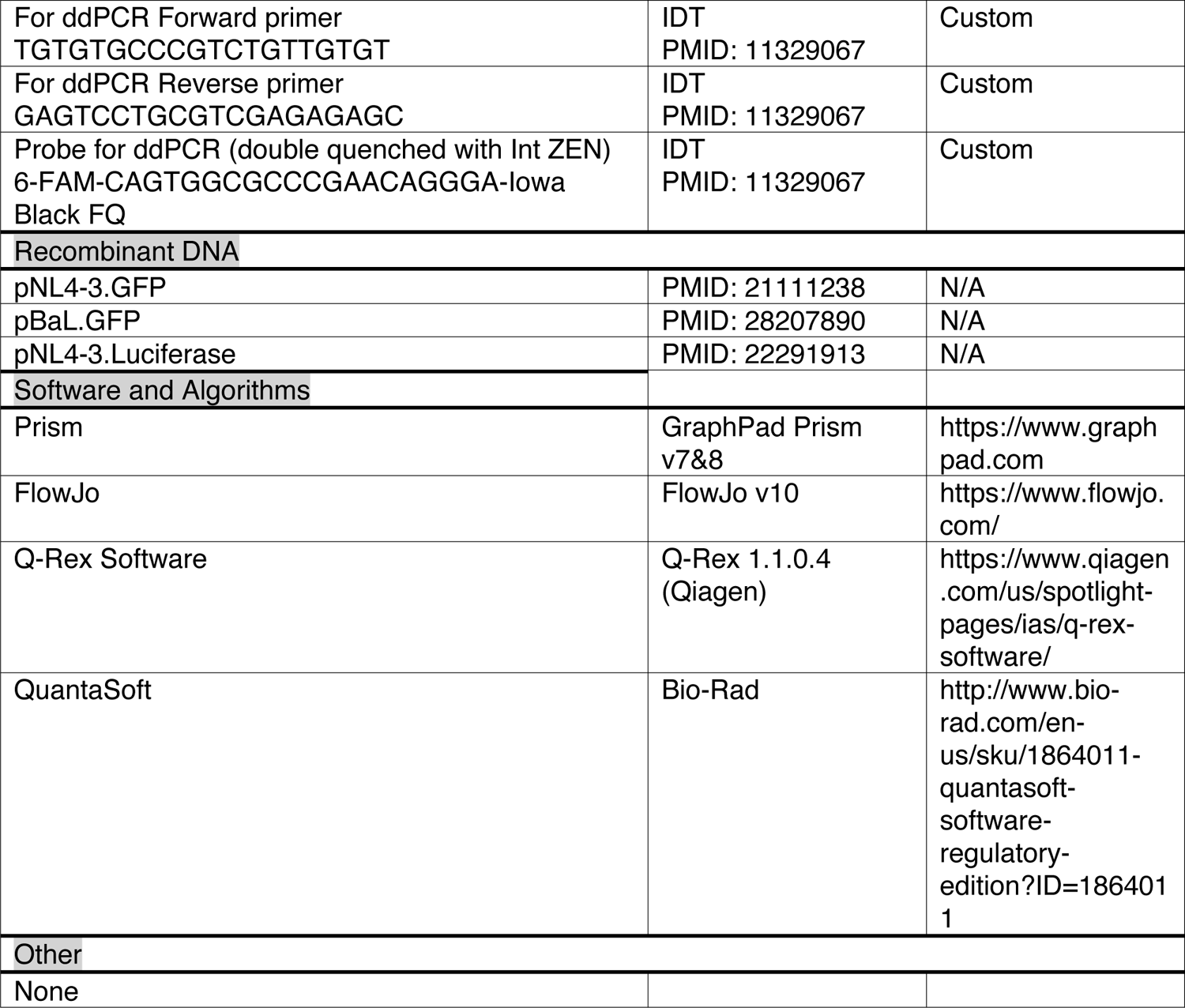

